# *Track-A-Worm 2.0*: A Software Suite for Quantifying Properties of *C. elegans* Locomotion, Bending, Sleep, and Action Potentials

**DOI:** 10.1101/2024.09.12.612524

**Authors:** Kiranmayi Vedantham, Longgang Niu, Ryan Ma, Liam Connelly, Anusha Nagella, Sijie Jason Wang, Zhao-Wen Wang

**Author notes:** **Correspondence** Zhao-Wen Wang, Ph. D.

## Abstract

Comparative analyses of locomotor behavior and cellular electrical properties between wild-type and mutant *C. elegans* are crucial for exploring the gene basis of behaviors and the underlying cellular mechanisms. Although many tools have been developed by research labs and companies, their application is often hindered by implementation difficulties or lack of features specifically suited for *C. elegans*. *Track-A-Worm 2.0* addresses these challenges with three key components: *WormTracker*, *SleepTracker*, and *Action Potential (AP) Analyzer*. *WormTracker* accurately quantifies a comprehensive set of locomotor and body bending metrics, reliably distinguish between the ventral and dorsal sides, continuously tracks the animal using a motorized stage, and seamlessly integrates external devices, such as a light source for optogenetic stimulation. *SleepTracker* detects and quantifies sleep-like behavior in freely moving animals. *AP Analyzer* assesses the resting membrane potential, afterhyperpolarization level, and various AP properties, including threshold, amplitude, mid-peak width, rise and decay times, and maximum and minimum slopes. Importantly, it addresses the challenge of AP threshold quantification posed by the absence of a pre-upstroke inflection point. *Track-A-Worm 2.0* is potentially a valuable tool for many *C. elegans* research labs due to its powerful functionality and ease of implementation.

## Introduction

The nematode *C. elegans* is commonly used as a model to probe the gene and neural circuit bases of behavior. Despite its small size and having only 959 somatic cells, *C. elegans* exhibits diverse behaviors, including locomotion, thermosensation [1], mechanosensation [2], chemosensation [3], social interactions [4], memory and learning [5, 6], sleep and arousal [7], and mating [8]. *C. elegans* also possesses many cellular properties observed in higher species, such as the generation of action potentials in muscle cells [9, 10] and some neurons [11, 12]. These behavioral and cellular properties are underpinned by a compact nervous system with only 302 neurons and a genome containing over 19,000 protein-coding genes [13], almost as many as the human genome (∼20,000) [14, 15]. Notably, more than 80% of *C. elegans* genes have human homologs [16], making it a particularly valuable model system for studying conserved gene functions.

All behaviors in *C. elegans* are dependent on locomotion. Consequently, mutations in various genes can lead to abnormal locomotor behaviors, such as changes in locomotor speed, forward and backward movement preference, and body bending and curvature. Therefore, analyzing locomotor behavior serves as a powerful method for elucidating the roles of genes in cellular and neural circuit functions. However, many mutant locomotor phenotypes are not easily detectable and quantifiable by the naked eye and require machine vision-based analysis. This need for machine vision has led to the development of automated worm tracker systems by numerous labs [17–33].

Existing worm trackers can be broadly categorized into two types: multi-worm trackers and single-worm trackers. Both types can be used to quantify mutant behavioral phenotypes. However, multi-worm trackers are essential for large-scale assays of pharmacological effects, whereas single-worm trackers are typically better suited for extracting detailed locomotion features. This is due to their higher magnification of the animal and their ability for long-duration recordings of the same worm using a motorized stage.

Most published automated worm-tracking systems were developed for in-house use and often had limited functionality. However, a few systems have been developed for public distribution. Notably, *WormLab* [34], a commercial multi-worm tracker from MBF Bioscience, is well-known. Another prominent system is the *Tierpsy Tracker*, an open-source multi-worm tracker [17, 18]. Both multi-worm trackers offer robust functionality but also come with various limitations. For instance, these limitations can include challenges in maintaining continuous tracking of the same worm due to the lack of a motorized stage, inability to differentiate the ventral and dorsal sides, and the quantification of numerous metrics that are not intuitive for typical *C. elegans* investigators. *Tierpsy Tracker* also has the added complexity of running the software in a containerized environment like Docker, which requires users to install and configure the software, manage dependencies, and address potential compatibility issues, presenting a steep learning curve for many researchers. Additionally, *Worm Tracker 2.0* [28], a single-worm tracker, was used to generate a behavioral database for many *C. elegans* mutants by the authors. Unfortunately, this previously open-source system is no longer available for download (https://worm-tracking.sourceforge.net/).

*C. elegans* undergoes periods of quiescence (approximately 2 hours each) between its developmental stages, which are referred to as developmentally timed sleep due to their resemblance to the sleep of other species in both behavior and molecular mechanisms. Machine vision is generally used to track the sleep state of worms restrained in microfluidic devices. However, restraining the animal within a tiny space might interfere with its sleep-like behavior. Therefore, a system capable of tracking freely moving worms for long durations would be highly useful for laboratories studying sleep-like behavior in *C. elegans*.

*C. elegans* body-wall muscle cells and some neurons can fire action potentials (APs) [9–12]. These APs typically result from calcium influx and, unlike those in mammalian neurons and muscle cells, lack a discernible inflection point in the rising phase, making automatic detection of the threshold challenging. As a result, investigators must determine the threshold either by manually inspecting each AP or by applying a fixed value across all APs in a recording trace. Both approaches have significant limitations, hindering the comparison of results across different research groups. The former is limited by subjective and potentially inconsistent criteria, while the latter does not account for baseline fluctuations. Since AP threshold is often used in quantifying AP amplitude, rise time, and afterhyperpolarization (AHP) amplitude, it is crucial to have software that enables an objective assessment of AP threshold in *C. elegans*.

*Track-A-Worm* is an open-source single-worm tracker based on the MATLAB. In its initial release [30], our goal was to provide the *C. elegans* research community with a system capable of quantifying locomotor and bending metrics useful to general *C. elegans* investigators while avoiding those that are primarily relevant to individuals with specialized computational expertise. Although this goal was essentially met, we have identified several significant limitations over the years. For instance, it does not quantify body length and curvatures, nor does it distinguish between the ventral and dorsal sides of the animal. Additionally, it cannot track worms on a bacterial lawn or control external devices for applications such as optogenetic stimulation. A significant software bug has also emerged with subsequent MATLAB updates, hindering accurate measurements of locomotion speed and the reconstruction of the worm’s travel path. Furthermore, a lack of video tutorials has made it challenging for readers to fully appreciate the system’s powerful functionality and ease of implementation.

Here, we present the release of the *Track-A-Worm 2.0* software suite, which consists of three components: *WormTracker*, *SleepTracker*, and *Action Potential (AP) Analyzer*. *WormTracker* has addressed all the issues of the original version identified above, with improved user interfaces and other new features. *SleepTracker* is used to record and analyze developmentally timed sleep in freely moving worms. *AP Analyzer* can detect the threshold using a definable and convenient approach and quantify other properties of action potentials in *C. elegans*. Importantly, video tutorials for each module of the system and analysis results of selected mutants are provided to guide users in using the system and to allow them to determine if it is the right solution for their lab.

### SYSTEM COMPONENTS

#### Software

*Track-A-Worm 2.0* software suite is available in two versions: a standalone version and a MATLAB-dependent version (https://health.uconn.edu/worm-lab/track-a-worm/). The standalone version can be used directly with the hardware described below, referred to as “standard hardware”. The MATLAB-dependent version is also configured for standard hardware but offers the flexibility to configure custom hardware (requiring basic MATLAB knowledge).

#### Hardware

##### Basic Hardware

1. A desktop or laptop computer running Windows 10 or 11 (64-bit) with a GigE (Gigabit Ethernet) port for the camera described below.
2. A stereomicroscope with an illuminated base. While most stereomicroscopes commonly used in *C. elegans* research laboratories are suitable for this application, those with good optics and a base that allows adjustable angled illumination offer advantages in image quality.

##### Additional Hardware for WormTracker

1. Motorized Stage: Optiscan^TM^ ES111, including a stage controller (ES11), a universal specimen holder (H473), a stage mounting bracket (H413), and a joystick (CS1521DP) (all from Prior Scientific, Rockland, MA, USA) (**Table 1**). The joystick is optional. You may want to request the manufacturer to reduce the stage mounting bracket height from the standard 8.0 cm to 2.0 –2.5 cm to allow better illumination from the microscope base. This stage has a travel range of 125 mm x 75 mm with a minimum step size of 1 µm. If you choose to use a different motorized stage, ensure that its travel distances (in both the *x* and *y* directions) are significantly longer than the diameter of the Petri dish used for worm tracking. Additionally, ensure that the stage position can be adjusted promptly so that adjustments can occur and complete between image acquisitions.
2. Camera: Mako G-040B (Allied Vision, Stadtroda, Germany). This black-and-white C-mount camera (**Table 1**) has a resolution of 728 x 544 pixels and uses GigE for video output. The relatively low resolution of this camera allows for smaller image file sizes and faster image processing. Images captured by this camera can be easily converted to high-quality binary images by the *WormTracker* software, even for those acquired in the presence of a bacterial lawn. This is in sharp contrast to the Sony XCD-V60 used in our original *WormTracker* [30], which prohibits tracking worms on a bacterial lawn. We strongly recommend using the Mako G-040B camera, though many newer cameras are likely to work equally well. To operate this camera, you will need either a dedicated power supply, such as the Allied Vision 12V 2A 8-pin Hirose Power Supply (**Table 1**), or a power over Ethernet (PoE) single port injector (**Table 1**). If you choose a different camera, ensure it allows precision time control by MATLAB and can be configured by MATLAB to a resolution of 728 x 544 pixels.
3. External Device Controller: myDAQ University Kit (National Instruments, Austin, TX, USA). This device (**Table 1**) is not required for typical worm tracking. *WormTracker* has been configured to control up to three external devices using myDAQ via TTL signals.
4. Fluorescence Light Source: To apply optogenetic stimuli during worm tracking, a fluorescence light source with adequate power and the ability to be controlled by TTL signals is required. Additionally, a longpass filter with a 520 nm cutoff is necessary in the illuminating light path to prevent blue light from desensitizing GCaMP6. In our setup with a Leica M165 FC fluorescence stereomicroscope, we place a 520-nm longpass cutoff filter (Y52, Hoya Corporation, Tokyo, Japan, **Table 1**) directly on the circular glass of the microscope base, which transmits light from a halogen bulb.

A photo of our *WormTracker* hardware system is shown in **Figure 1**.

**Figure 1.**
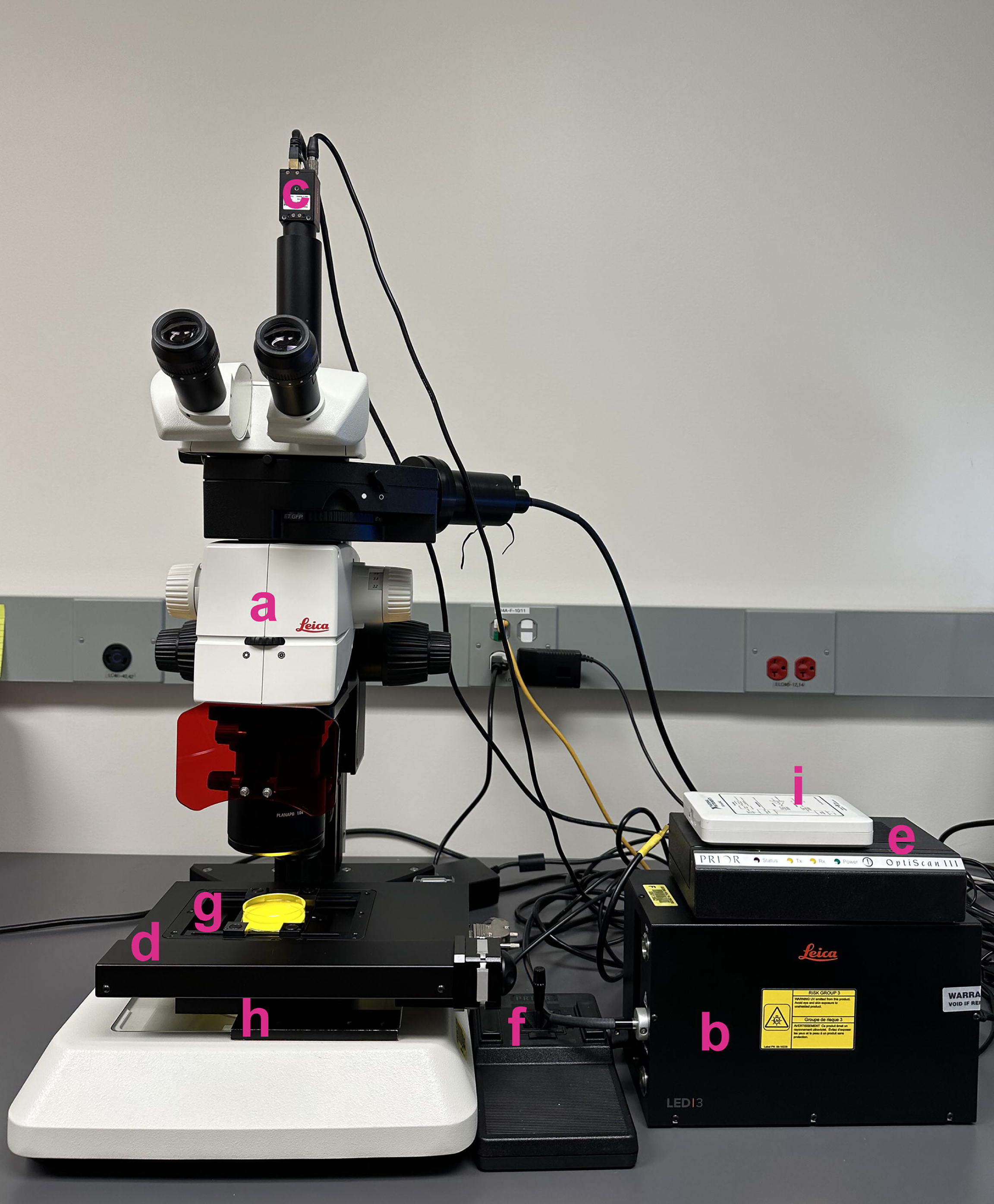
*WormTracker* hardware components. Shown are the hardware configuration of our system, including a fluorescence stereomicroscope (M165 FC, Leica) (a) with a fluorescence light source (LED3, Leica) (b), a C-mount CMOS camera (Mako G-040B, Allied Vision) (c), a motorized stage (Optiscan^TM^ ES111) (d) with a stage controller (ES11) (e), a joystick (CS1521DP) (f), a universal specimen holder (H473) (g), and a stage mounting bracket (H413) (h) (all from Prior Scientific), and an external device controller (myDAQ, National Instruments) (i). The *WormTracker* software is preconfigured to work with components c, d, and i. Components b and a 520-nm longpass cutoff filter (Y52, Hoya Corporation) are required only for optogenetic stimulation. The Petri dish is 60 mm in diameter. The yellow color in the Petri dish is due to light from the microscope base filtered through the longpass filter.

##### Additional Hardware for SleepTracker

1. Camera: DMK 37BUX273 (The Imaging Source, Bremen, Germany). This black-and-white C-mount camera (**Table 1**) has a resolution of 1440 x 1080 pixels and uses USB 3.1 for video output. Due to its higher resolution, we use this camera instead of the Mako G-040B for tracking worm sleep behavior.
2. Sleep Recording “Chamber”: We use a rectangular polydimethylsiloxane (PDMS) membrane (14.0 mm x 10.5 mm x 0.65 mm) containing six circular apertures (3.0 mm in diameter) for recording sleep behavior. This membrane forms six “chambers” when its bottom is in contact with the agarose nematode culture plate. A CAD design of the sleep chamber is provided in **Appendix A**.

### TRACKING AND ANALYSIS ALGORITHM

#### WormTracker

##### 1. Stage Tracking

To record a worm’s locomotor activity at high resolution, it must be kept near the center of the camera’s imaging field using a motorized stage. *WormTracker* compares the worm’s positions across successive images and adjust the stage position accordingly at 1-second intervals. All stage movements are automatically recorded in a stage file, which is later used in combination with worm image files for analyses. Stage positions are corrected between image acquisitions to prevent blurring and discontinuities in the acquired images.

##### 2. Ventral and Dorsal Differentiation

Mutations in certain genes may alter body bending with a ventral or dorsal bias. *WormTracker* can quantify this bias through body curvature analysis with ventral and dorsal distinction. However, since automatic differentiation of the ventral and dorsal sides is essentially impossible, the user must visually determine the orientation through the microscope eyepieces and enter the ventral/dorsal information in the *Record* module before starting the recording.

##### 3. Image processing

Recorded images are processed through several steps to produce a spline. The original video frames are grayscale images, with pixels having brightness values between 0 (black) and 255 (white). Each video frame is converted into a binary image using a user-selected brightness threshold, such as 100. Pixels with brightness values above the threshold are converted to pure white, while those below it are converted to pure black. If error-checking algorithms flag the video frame as unanalyzable, the software automatically adjusts the threshold by −20 to +20% over up to 10 attempts. In the binary image, all objects smaller than 3000 pixels in area (0.76% of a 728 × 544 pixel image) are removed to avoid complications from debris and other artifacts, and any gaps caused by light spots within the worm’s body are filled in. An edge detection algorithm is then applied to find the worm’s outline, and the raw edge data are smoothed and converted into a series of x-y coordinates. These procedures transform a grayscale image into a binary image, where the worm appears as a solid black shape (**Figure 2a**).

**Figure 2.**
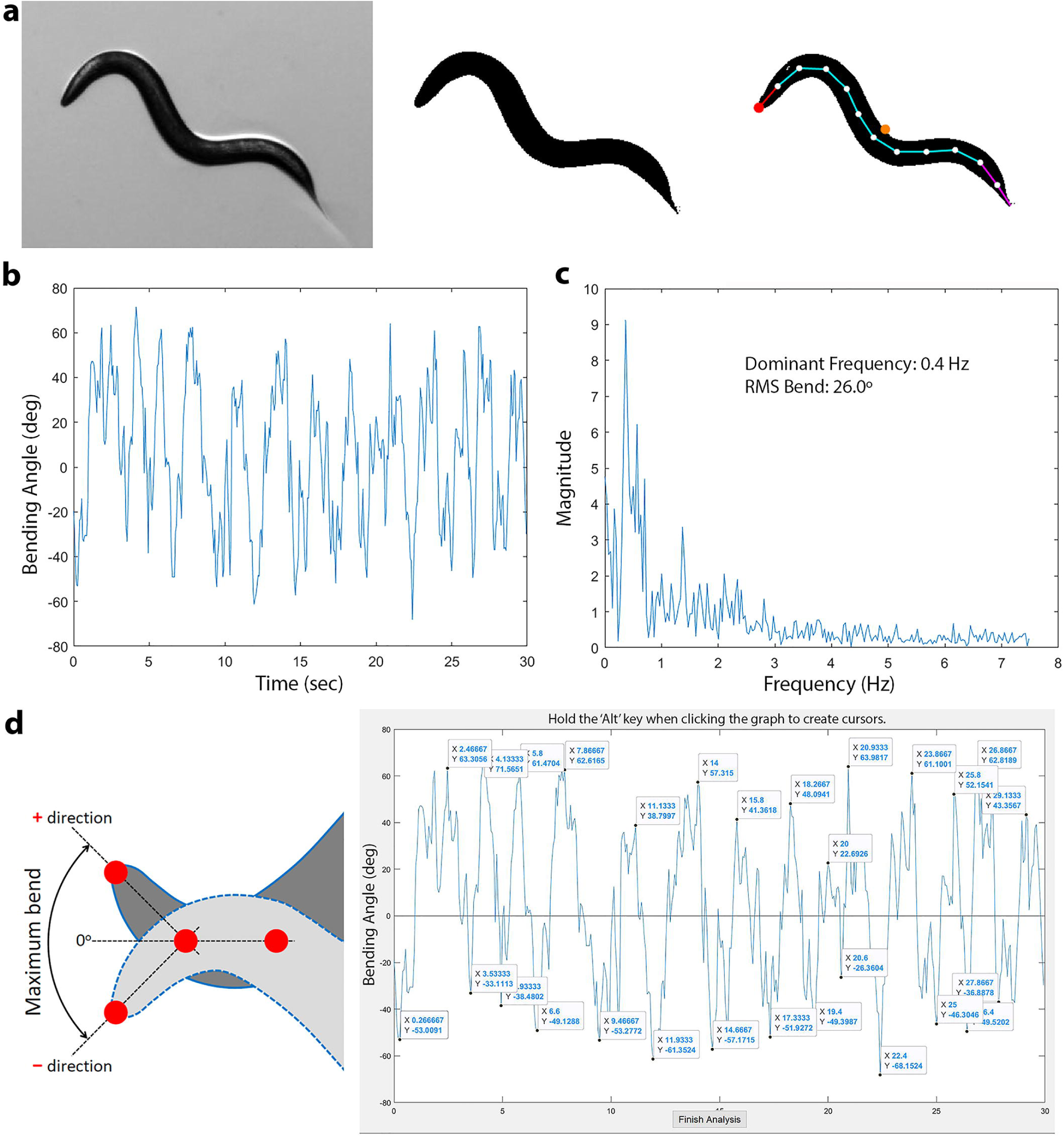
Spline generation and bending property quantification. **a.** Conversion of a grayscale image (*left*) to a binary image (*middle*), followed by spline fitting to the binary image (*right*). The spline is generated by placing 13 markers at equal intervals along the longitudinal axis of the worm’s body, with the marker at the head tip shown in red and the centroid in orange. **b.** Bend trace of a worm recorded over 30 seconds. **c.** Bending frequency spectrum of the same worm, with the dominant bending frequency and RMS (root mean square) bend angle from the Results section displayed within the graph. **d.** Quantification of the maximum bend. *Left:* Diagram illustrating the definition of the maximum bend, which is the maximum bending angle of a spline marker point in the ventral and dorsal directions relative to a straight line fitted to the two subsequent spline markers. *Right:* A bend trace with *x* and *y* coordinates marking the times and bending angles of alternative peaks and troughs. The coordinates are determined by clicking on the peaks and troughs, using a 20-degree threshold.

##### 4. Head Identification

In the binary image, the tail of the worm forms a sharper angle than the head. A custom corner-detection algorithm is used to identify the sharpest and second-sharpest corners to designate them as the tail and head candidates, respectively. This approach is similar to that used in a previous study [35]. Error-checking mechanisms are incorporated into this process to ensure that corners formed by body bends are not mistaken for the head or tail. Head identification in each subsequent frame also uses the head position from the preceding frame as a reference to facilitate the process.

##### 5. Spline Generation and Centroid Determination

A spline is generated along the midline of the worm profile by performing a cubic interpolation of midpoints (**Figure 2a**). This spline is divided into 12 segments by placing 13 markers at equal intervals, with the centroid defined as the average position of the 13 markers.

With decent image quality and an appropriate threshold for spline fitting, *WormTracker* can automatically produce accurate splines with a 100% success rate, except when the worm is in a coiled shape. Typically, the software successfully fits the spline on the first attempt. However, a non-optimal fitting threshold, poor image quality, or a coiled worm body shape (e.g. an Ω shape) can hinder spline fitting. In such cases, the software automatically adjusts the brightness threshold by up to 20% in either direction and repeats the analysis until an acceptable spline is extracted or a total of 10 attempts have been made. Images that cannot be successfully analyzed after 10 attempts are set aside for semi-automatic analysis.

##### 6. Bending Property Quantification

###### WormTracke

plots the bending activity of a worm over the recording period as a Bend Trace (**Figure 2b**). The appearance of a bend trace provides the user with a general overview of the worm’s bending properties. The most important use of the bend trace is to serve as the basis for detailed quantifications of bending properties, including Dominant Bend Frequency, Maximum Bend, and RMS (root mean square) bend. The bending properties at each spline marker point are quantified using the reference of a straight line connecting the two subsequent markers. For example, the bending activity at marker 1 uses the straight-line fit of markers 2 and 3 as the reference, while that at marker 11 uses the straight-line fit of markers 12 and 13. As a result, bending activities can be quantified for markers 1 through 11, which we sometimes refer to as “bends” in the text.

###### Dominant Bend Frequency

A worm typically bends with one large dominant motion interspersed with smaller oscillations. A Fourier transform of the bend trace is performed to obtain the frequency spectrum (**Figure 2c**), where the most prominent frequency peak is defined as the dominant bend frequency.

###### Maximum Bend

The Maximum Bend is calculated as the difference between the averages of the negative and positive values of the dominant bends in a bend trace (**Figure 2d**). The dominant bends represent the most ventral and dorsal bending angles. This measure is particularly useful for detecting organized mutant bending behavior but has limited value for identifying chaotic bending behavior without regular large-amplitude bends. Its usefulness is demonstrated by the significant increase in Maximum Bend at marker 1 in loss-of-function mutants of *slo-1*, which encodes the BK channel (Slo1) in *C. elegans*, compared to the wild type [36].

###### RMS Bend

While *Maximum Bend* is an intuitive measure of bending angles, it does not capture all aspects of bending property. A bend trace often includes numerous smaller oscillations that are not accounted for in the analysis of maximum bends (**Figure 2b**). *RMS Bend* provides an objective, sensitive, and accurate measure of overall bending activity, calculated using the following formula:

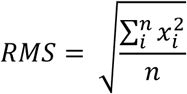

where *n* is the total number of data points, and *i* refers to the index of each individual point. While RMS Bends have both positive and negative values, as seen in the exported Excel file, the Results section of the *Analysis* module displays an average value of RMS Bends based on absolute values. However, RMS Bend is limited in that it does not distinguish between low-amplitude, high- requency, and high-amplitude, low-frequency bending behavior.

###### Sum of All Bends

The *Sum of All Bends* is a useful metric for quantifying the bending behavior of an entire worm. It is calculated as the sum of all 11 RMS Bends.

##### 7. Speed, Distance, and Direction Quantification

These metrics are typically based on the positions of the centroid over successive images, although any of the 13 markers along the spline may be used. The calculation of these movement metrics is based on the actual frames if the frame rates are 1, 3, or 5 per second, but defaults to 3 frames per second if the recording was acquired at 15 frames per second. We chose 3 frames per second as the default because it is more than adequate for quantifying movement metrics.

*WormTracker* measures the average speed (µm/sec), the total distance traveled (both forward and backward), as well as the net distance traveled (the straight-line distance between the first and last positions of the worm) over the recording period. Directionality is determined by comparing the velocity vector (from the last centroid to the current centroid) with the head vector (from the current centroid to the current “nose” / head tip). If the projection of the head vector onto the velocity vector is positive, the worm is determined to be moving forward, and vice versa (**Figure 3a**).

**Figure 3.**
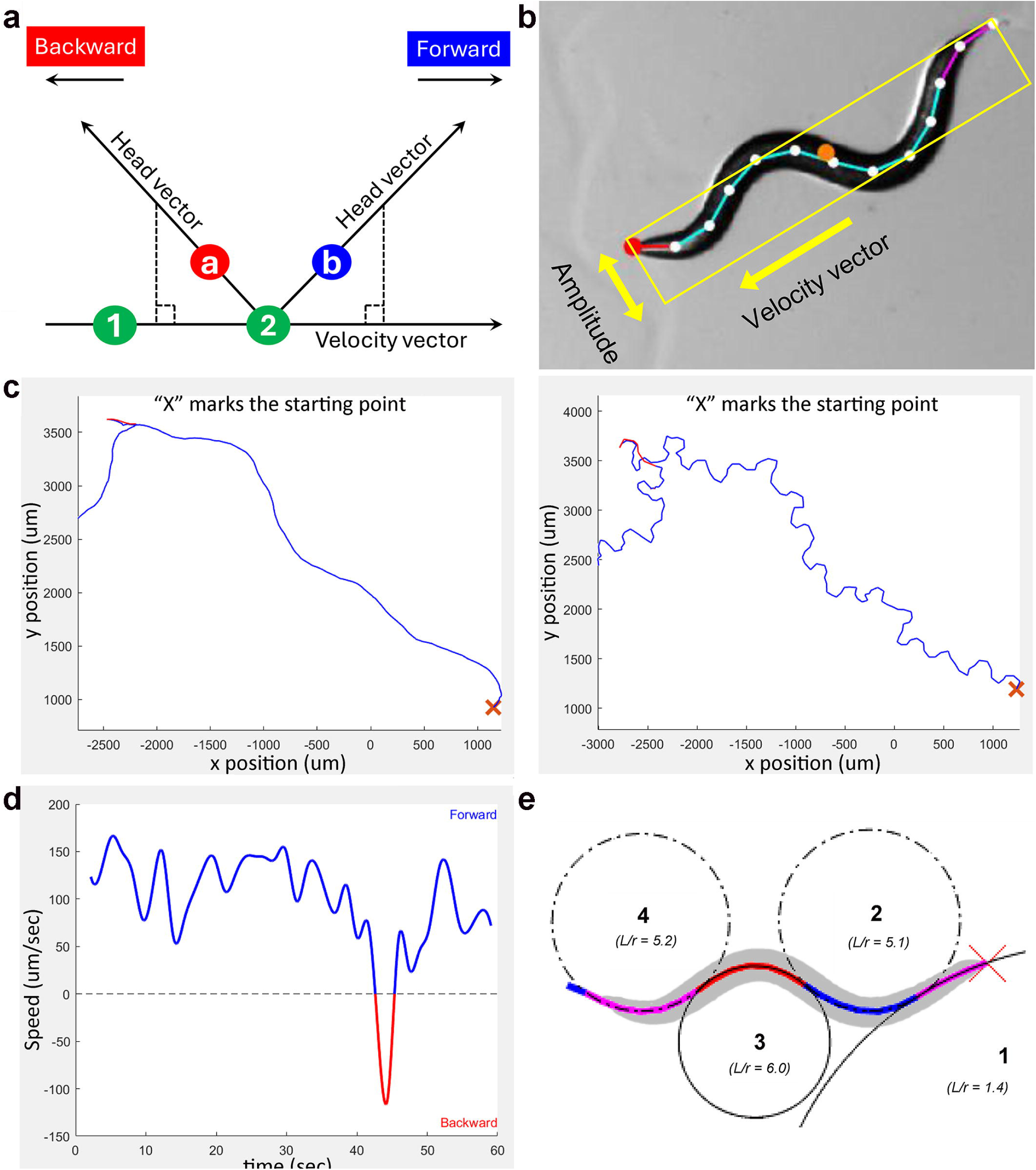
Analysis of locomotion directionality, speed, and body curvatures.a. Directionality is determined by examining the relationship between the velocity vector and the head vector. The velocity vector connects the centroids of the current (#2) and previous (#1) frames, while the head vector connects the current centroid to the current head point (a or b). If the head vector projects onto the positive side of the velocity vector, the worm is moving forward; otherwise, it is moving backward. **b.** Method for quantifying worm amplitude. A rectangle is drawn parallel to the velocity vector, just large enough to enclose all spline markers. The width of the rectangle is the worm’s amplitude. **c.** Reconstructed worm travel paths based on the positions of the centroid (*left*) and spline marker 1 (*right*). **d.** Plots of forward and backward locomotion speeds over time shown for interpolated data. **e.** Method for quantifying body curvatures. Circles are fitted to worm segments, with curvature values calculated as the worm length (*L*) divided by the radius (*r*) of each fitted circle (*L/r*).

##### 8. Length and Amplitude

The length of the worm is determined by measuring the length of the straightened spline. Amplitude is a measure of worm curvature. It is determined by first finding the velocity vector as described above and then measuring the width of a rectangular box aligned with the velocity vector and just large enough to contain all spline points (**Figure 3b**). The software also calculates the ratio of amplitude over worm length (A/L) to account for the effects of worm size variation on the measurement. The worm length (L) here refers to the actual worm length, not the length of the rectangular box.

##### 9. Travel Path Reconstruction

Displaying a worm’s travel path in publications or presentations is often useful. *WormTracker* can reconstruct this path by integrating consecutive frames of worm images with information from the corresponding stage and time files. This travel path can be generated based on the positions of the centroid or any of the 13 markers along the spline (**Figure 3c**). For recordings at 1, 3, or 5 frames per second, the worm path is plotted according to the actual frame rate. For recordings acquired at 15 frames per second, the worm path is plotted at 3 frames per second.

*WormTracker* also reports the distances traveled and speeds during forward and backward movement, and generates a plot of forward/backward speed over time using either binned or unbinned data. The plot can be based on either raw or interpolated data (**Figure 3d**), with the latter used to smooth out the trace.

##### 10. Body Curvature Quantification

The *Curve Analyzer* module first generates a smooth curve along the 13 spline markers and then identifies its inflection points. Short segments, defined as shorter than 3% of the spline length, are excluded to remove artifacts. Circles are then fit to the worm segments using the Kasa method, a least-squares algorithm that finds the best-fit circle by minimizing the squared distance between the points and the circle’s perimeter. Curvature values are calculated as the worm length divided by the radius of each fitted circle (L/r) (**Figure 3e**). The normalized midpoint of each segment is computed by averaging the start and end positions of the segment and then dividing by the total worm length.

#### SleepTracker

*SleepTracker* detects sleep-like behavior based on worm locomotor activity, which is measured by quantifying the differences of centroid locations between consecutive images. Typically, worm images are acquired at a rate of 1 frame every 10 seconds. During analysis, the worm is considered to be in a motionless state if the difference between centroid locations of two consecutive frames is <10 µm, and in an active state if the difference is >10 µm. For each recording, the beginning of the first three consecutive frames of motionless state marks the time point of entering sleep, while the end of the last three consecutive frames of motionless state marks the time point of exiting sleep. Total sleep duration is defined as the time from entering to exiting sleep (**Figure 4**). The use of third-party software (details provided in a subsequent section) also allows for the quantification of active events during sleep and the duration of motionless sleep, which is calculated by subtracting the summed duration of active events from the total sleep duration.

**Figure 4.**
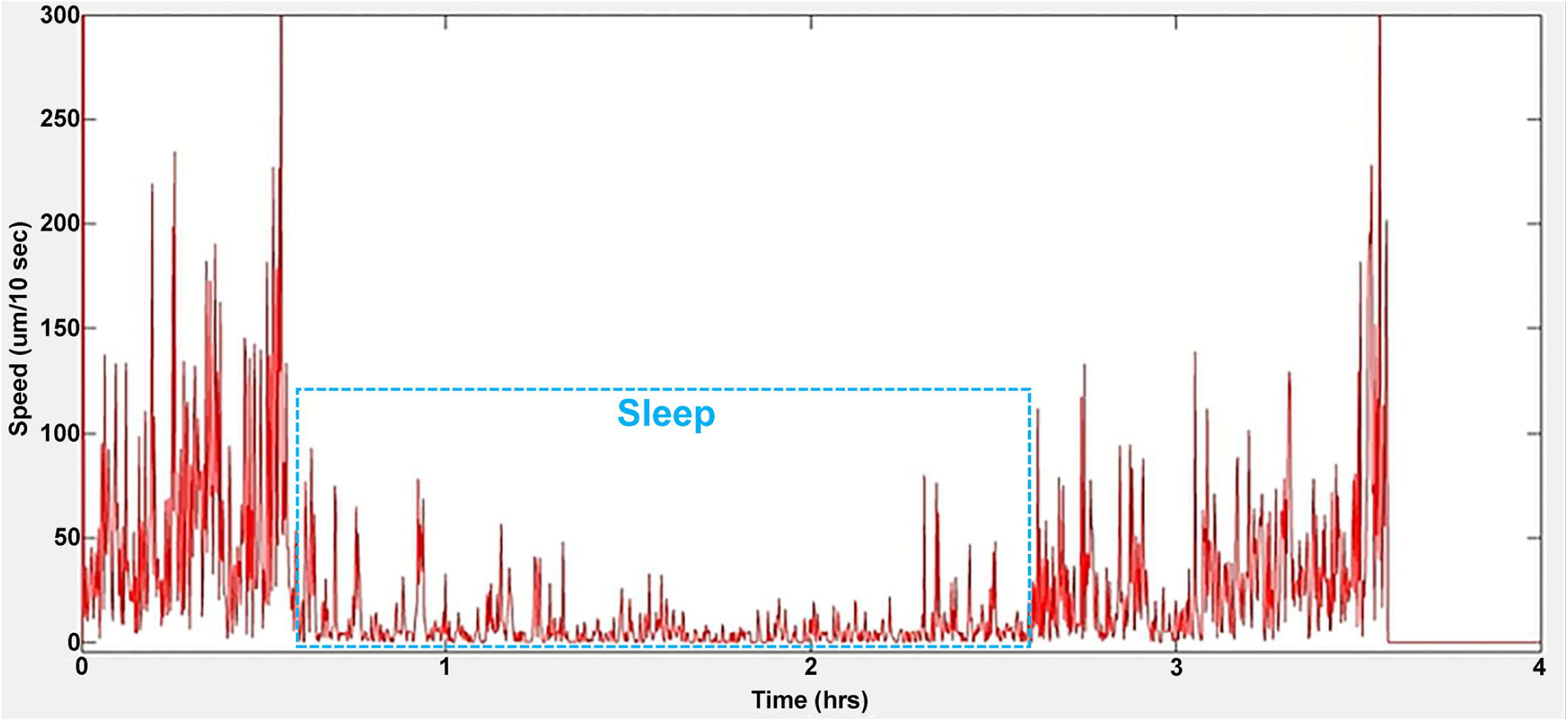
Quantification of sleep duration in freely moving worms using *SleepTracker*. The actogram shows the locomotor activity of a worm developing from the L4 to young adult stages. Images were captured at a rate of 1 frame every 10 seconds. Sleep duration, highlighted by the rectangular box, is defined as the period from the start of three consecutive motionless frames to the end of the last three motionless frames. The worm is considered motionless if the difference between the centroid positions of two consecutive frames is <10 µm. Spikes within the sleep period are classified as active events.

#### AP Analyzer

Nearly all electrophysiological experiments with *C. elegans* are conducted using the *pClamp* acquisition system (Molecular Devices, San Jose, CA). However, *ClampFit (version 11)*, the analysis module of this system, does not offer automatic detection of AP thresholds, requiring users to manually select a membrane voltage as the threshold for all APs being analyzed. *AP Analyzer* addresses this by allowing users to import AP data obtained with *pClamp* and identify AP thresholds using two different criteria, depending on whether an inflection point is present before the AP upstroke. For APs without an inflection point, such as those in *C. elegans* body-wall muscle cells (**Figure 5a**), users must manually select a specific time before the AP peak, and this time point is then used to determine the threshold for all APs. Although this pre-peak time selection is still subjective, applying it consistently across all APs ensures reproducible assessment of threshold values, as well as associate AP rise times and AP amplitudes. For APs with an inflection point preceding the upstroke (**Figure 5a**), the threshold is detected automatically.

**Figure 5.**
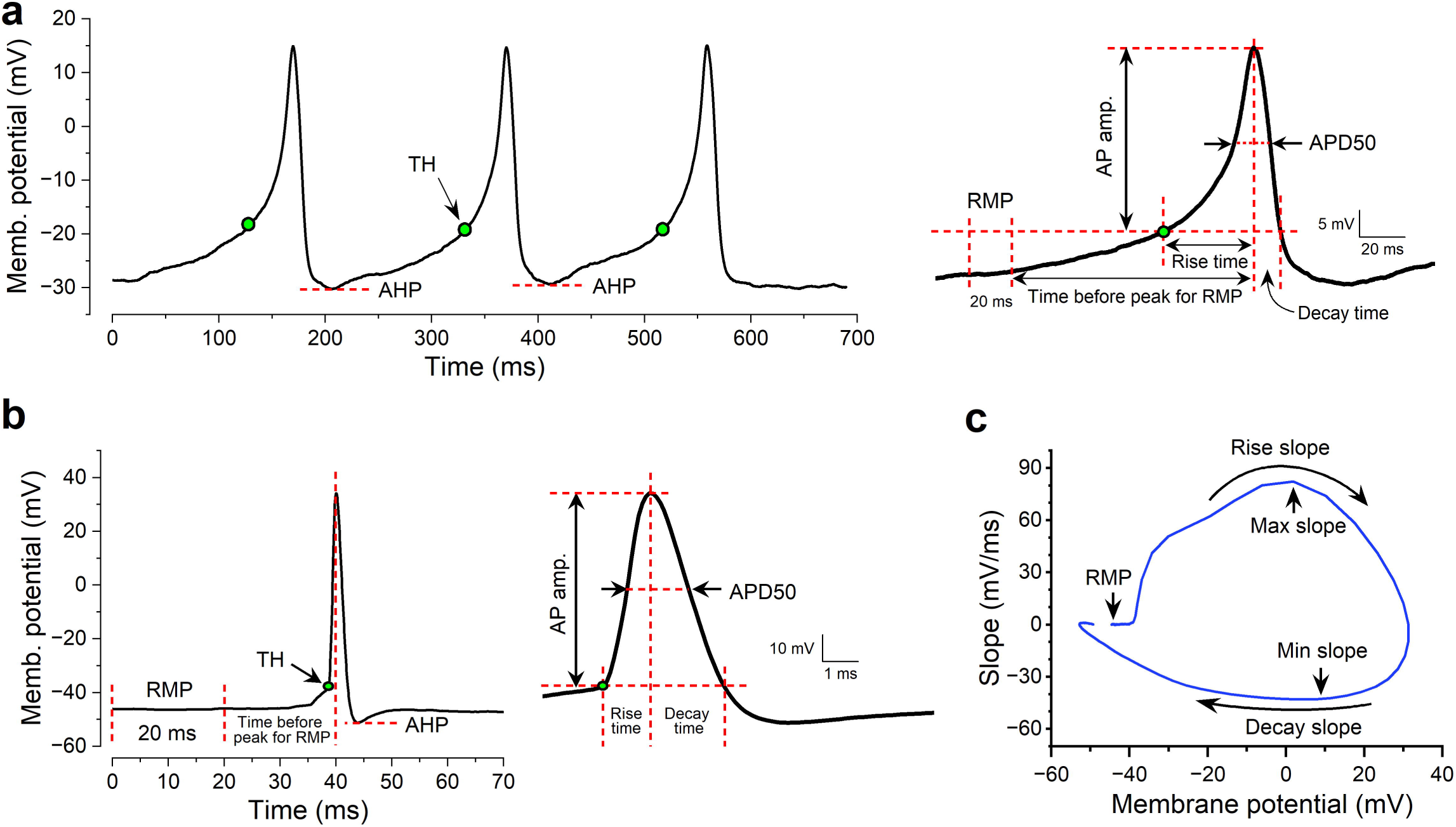
Quantification of action potential (AP) properties using *AP Analyzer* with two different approaches. **a.** AP threshold is the membrane potential at a user-defined pre-AP peak time for APs lacking a pre-upstroke inflection point. **b.** AP threshold is automatically detected for APs with a pre-upstroke inflection point. The representative APs are from a body-wall muscle cell of a wild-type *C. elegans* (**a**) and a suprachiasmatic nucleus (SCN) neuron of a wild-type CBA/CaJ mouse (**b**). In both cases, the resting membrane potential is calculated as the average membrane potential over 20 ms before a user-selected pre-AP peak time. The rise time is defined as the time from AP threshold to AP peak, the decay time from AP peak back to the membrane potential matching the AP threshold, and afterhyperpolarization (AHP) as the actual membrane voltage. **c.** Voltage phase plot of an SCN neuron AP from a mouse, illustrating the quantification of AP maximum and minimum slopes, corresponding to the points where the membrane potential increases and decreases most rapidly, respectively.

The quantified results include values for resting membrane potential (RMP), AP threshold, AP amplitude, APD50 (AP duration at 50% amplitude), AHP level, AP rise and decay times, and the maximum and minimum slopes of the AP. The rise time refers to the time taken to depolarize from the threshold to the AP peak, while the decay time represents the time taken to repolarize from the AP peak back to the threshold level (**Figure 5a, b**). The maximum and minimum slopes are derived from a voltage phase plot generated by the software (**Figure 5c**). The software calculates the RMP as the average membrane potential over a 20 ms period before a user-selected pre-AP peak time. This RMP value is accurate if inter-spike intervals are long, but it may deviate from the true RMP if inter-spike intervals are short, preventing the membrane potential from fully returning to the true RMP. In such cases, users should manually quantify the RMP from a segment of the recording trace with few or no APs using *ClampFit*.

#### WormTracker File Organization

WormTracker relies on a specific file structure to function properly. It integrates data from various files, such as image files, stage files, time files, and spline files, which must be organized hierarchically for automatic access. Adhering to this structure is essential, and all files should be saved in their designated folders. To clarify the file organization, refer to Figure 4, which illustrates the hierarchical structure of the *WormTracker* files.

#### Example: Recording 10 Wild-Type Worms

Suppose you’re recording 10 wild-type worms, naming each worm from wt1 to wt10, and saving them in a parent folder named “Wild Type.” Follow these steps:

1. Create Folders: In Windows Explorer, create the folder structure Wild Type\wt1\.
2. Record Images: In the Record module, select the wt1 subfolder in the Image Folder section and record the first worm’s images.
3. Update Folder: Change “1” to “2” in the Image Folder path, press Enter, and record images of the second worm in the updated wt2 subfolder.
4. Repeat: Follow Step 3 for the remaining 8 worms.

#### Recording File Organization

After recording, the “Wild Type” parent folder will contain 10 subfolders (wt1 to wt10), each holding the respective worm’s image files (e.g., img00001.jpeg, img00002.jpeg). If you recorded with ventral and dorsal distinction, the file names will have an “R_” or “L_” prefix, indicating the ventral side was on the right or left.

Each worm’s subfolder (wt1, wt2, etc.) will also contain a stage file (wt1.txt) and a time file (wt1_times.txt). If the frame rate was adjusted using the Playback module, a “Removed” subfolder will be created with the discarded images, and a new time file will replace the original, which will be renamed (wt1_times_before_RF.txt). The Restore tab in the Playback module can revert all changes in the subfolder.

#### Spline Fitting File Management

When fitting splines for images in the wt1 subfolder using the Fit Spline or Batch Spline module, the resulting spline file (wt1_spline.txt) is saved in the “Wild Type” parent folder, not the subfolder.

By following these procedures, WormTracker will:

1. Automatically identify the spline file in the Fit Spline module when loading a recording, if it exists.
2. Automatically identify the spline file in the Playback module when loading a recording, if it exists.
3. Automatically load the stage and time files in the Analyze and Batch Analyze modules when loading a spline file.

*Recommendation:* Save Excel analysis files in the same “Wild Type” parent folder for easy access.

### SOFTWARE INTERFACE OVERVIEW AND FUNCTIONALITY

*Track-A-Worm 2.0* includes multiple user software interfaces. The use of these interfaces is demonstrated in the video tutorials on your website (https://health.uconn.edu/worm-lab/track-a-worm/) with video captions in **Appendix B**. Below is an overview of the software interfaces and a description of their functionality.

#### WormTracker

*WormTracker* includes eight different modules: Calibrate, Record, Playback, Fit Spline, Analyze, Batch Spline, Batch Analyze, and Curve Analyzer. Each module has its own graphic user interface and may be launched from a common launch pad (**Figure 6a**).

**Figure 6.**
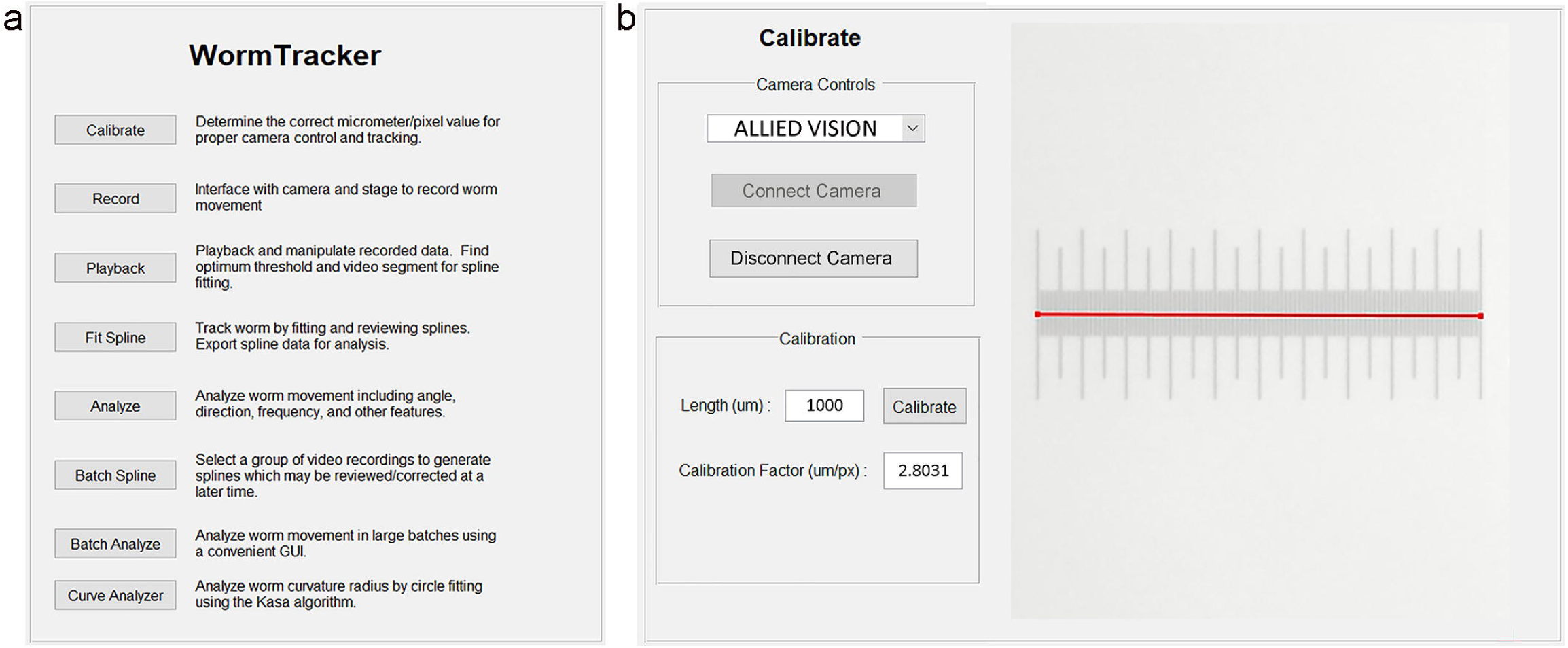
User interfaces of *WormTracker* launch platform and WormTracker Calibrate module. a. The launch platform allows users to start various *WormTracker* modules. b. The Calibrate module calibrates the camera’s pixel size to micrometers. To use it, place a micrometer in the camera’s field of view, click on both ends of the micrometer, enter its length, and click “Calibrate”. The resulting calibration factor shows the numeric relationship between micrometers and pixels.

#### Calibrate

Before recording images, it is essential to determine the correct conversion factor from pixels to micrometers using the *Calibrate* module (**Figure 6b**,). This factor must be entered into the *Record* module for accurate stage tracking.

#### Record

The *Record* module (**Figure 7**) receives worm position data from the camera, issues commands to re-center the stage at 1-sec intervals, and outputs sequential images to a user-designated folder. It also saves a text file containing stage movement information, and another text file containing the timing information for each captured image in the recording. The user can define the frame rate (1, 3, or 15/sec), recording duration, and the worm’s ventral/dorsal orientation. Up to three external devices can be controlled by TTL signals using any on-and-off patterns, and each device can be managed independently. The *Record* module can display the worm as either grayscale or binary images during recording and save the grayscale images at either 50% or 100% of the camera’s resolution. Before the recording, it is important to specify the file folder and the file name, center the stage, and enter the correct calibration factor obtained with the *Calibrate* module.

**Figure 7.**
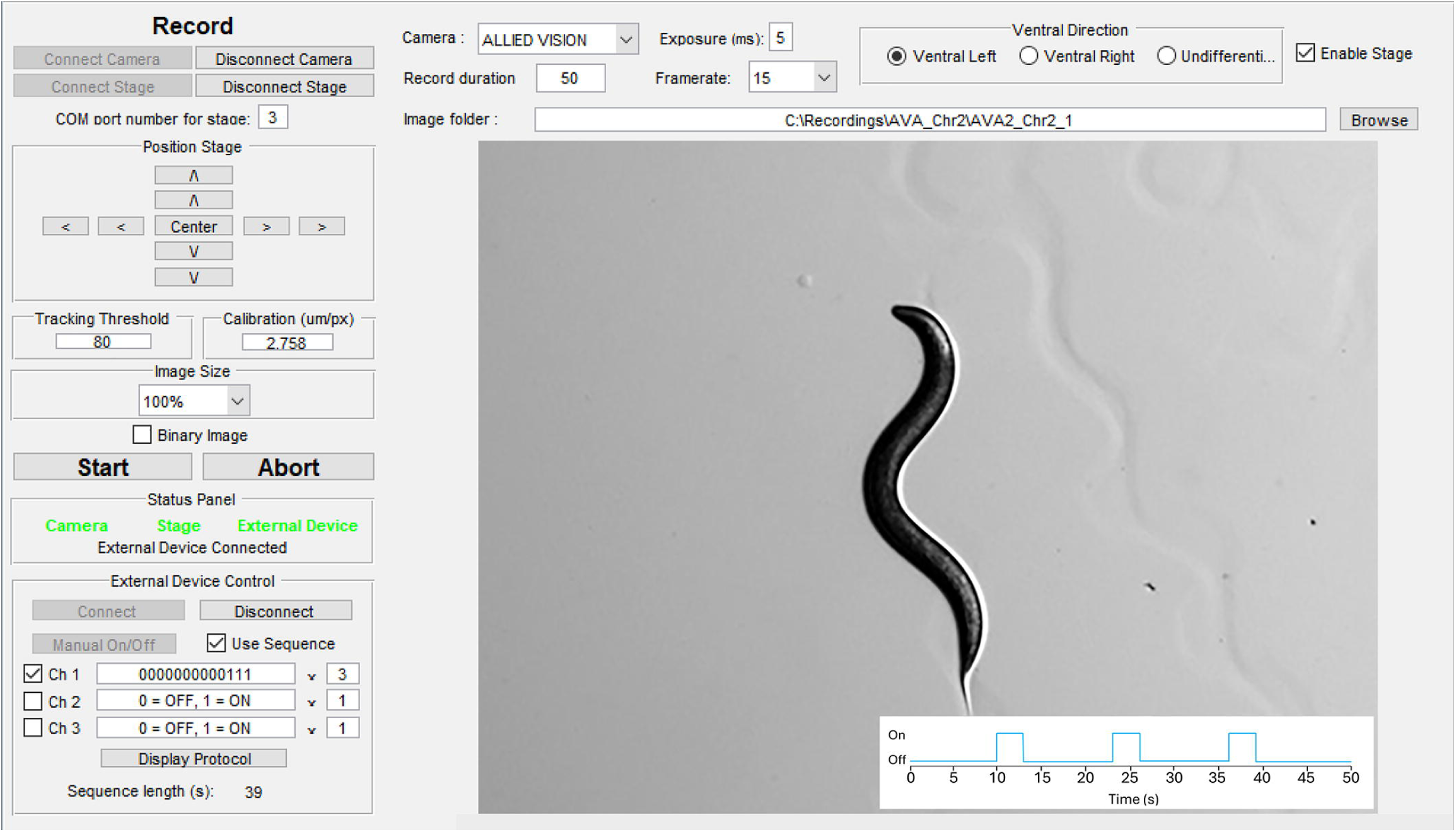
User interface of *WormTracker* Record module. This module captures grayscale snapshots of a freely moving worm at user-defined frame rates and recording durations. It keeps the worm near the center of the camera’s field of view using a motorized stage, allows the user to input ventral and dorsal orientation information, displays the worm in either grayscale or binary images, and can control up to three external devices via TTL signals. The camera imaging field shows a sample stimulus protocol, which matches the parameters defined for Channel 1 in this figure and is normally displayed in a separate window. Note that the Off period after the third stimulus is the difference between the recording duration (50 seconds) and the total stimulus protocol duration (13 seconds x 3 runs = 36 seconds).

#### Playback

The *Playback* module (**Figure 8**) allows users to view recorded sequential images as a movie, adjust the frame rate (by reducing or restoring it), and create movies from the recordings. Images can be displayed as either grayscale or binary format, with or without the fitted spline. If spline fitting has been performed for the recording, the corresponding spline file is automatically detected and can be activated by clicking the Load tab next to it. Movies can also be created in grayscale or binary, with or without the fitted spline. This module is particularly useful for initially assessing the threshold for spline fitting, quickly verifying spline fitting results, reducing frame rates for faster analysis, and creating movies for publications or presentations.

**Figure 8.**
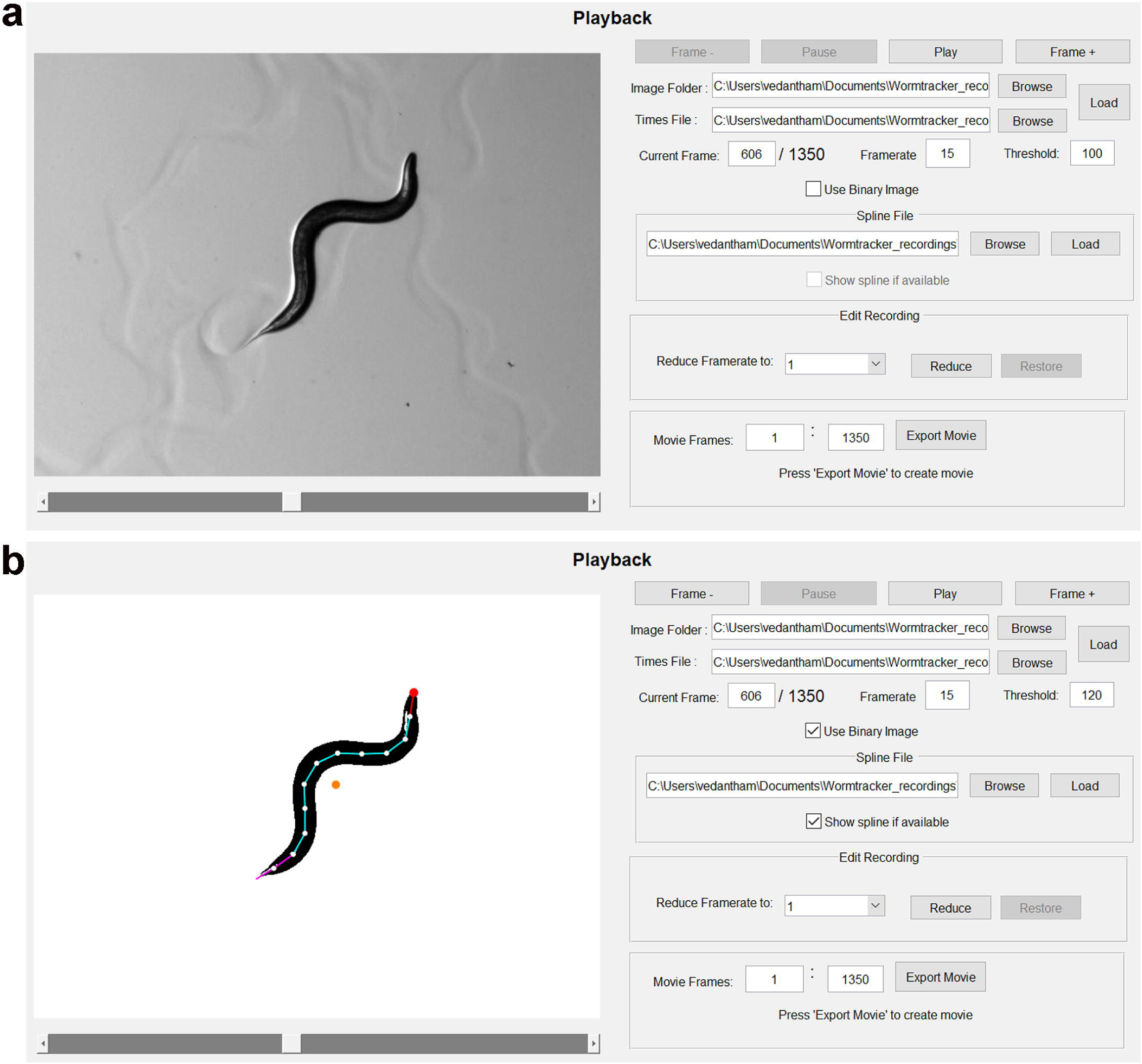
User interface of the Playback module. This module plays movies that show the worm as either grayscale (**a**) or binary (**b**) images, and without (**a**) or with (**b**) the fitted spline. It allows users to evaluate thresholds for converting pixels into black and while dots for spline fitting, adjust frame rates, and create movies (grayscale or binary, with or without the fitted spline) from recordings. The orange circle in (**b**) represents the centroid.

#### Fit Spline

This module (**Figure 9a and b**) has two functions: automatic head identification and spline fitting, and user-assisted verification and correction of the automatically generated spline results. The second function can be applied to splines generated automatically by either the *Fit Spline* or the *Batch Spline* module.

**Figure 9.**
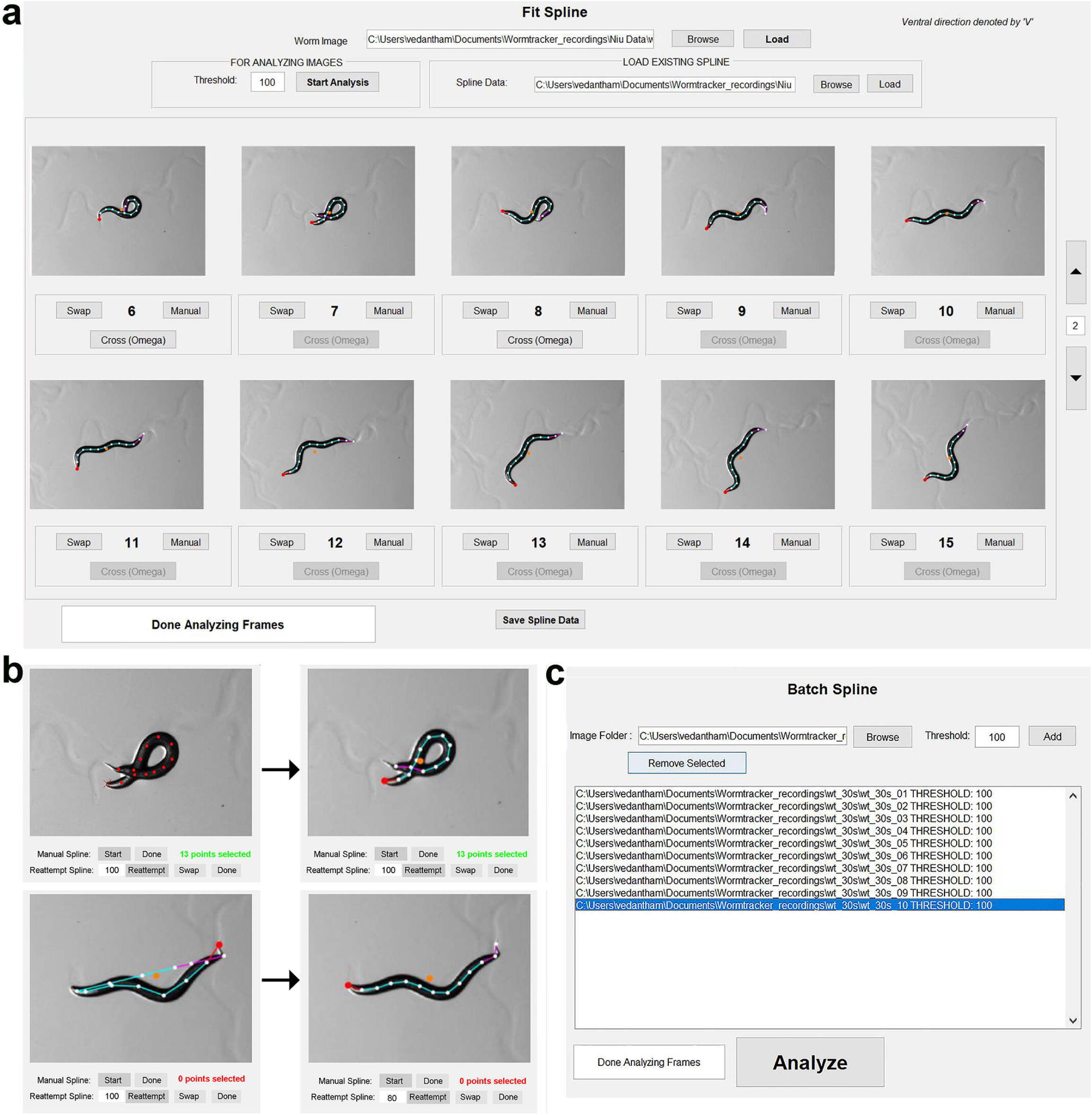
User interfaces of the Fit Spline and Batch Spline modules. **a.** User interface of the Fit Spline module. This module is used to fit splines to worm images based on a user-chosen threshold for binary conversion, and to verify and correct the fitting results. Shown are 10 successfully fitted frames, where the red dot represents the first spline marker, and the orange dot represents the centroid. **b.** Examples of spline fitting correction for specific frames using two different methods: (1) manually clicking along the worm 10 or more times from head to tail (top), and (2) redoing automatic spline fitting using a different threshold. **c.** User interface of the Batch Spline module. This module allows users to select multiple recordings for automatic spline fitting at the same or different thresholds.

#### Batch Spline

This module (**Figure 9c**) is designed for automatic spline fitting across multiple recordings, saving users the time spent waiting for results. However, like the results generated automatically by the *Fit Spline* module, the results from the *Batch Spline* module still require user-assisted verification and correction using the *Fit Spline* module.

#### Analyze

This module (**Figure 10a**) extracts quantitative and graphical information from a spline file and the corresponding stage and time files. It is divided into two sections: Bend Analysis and Movement Analysis.

**Figure 10.**
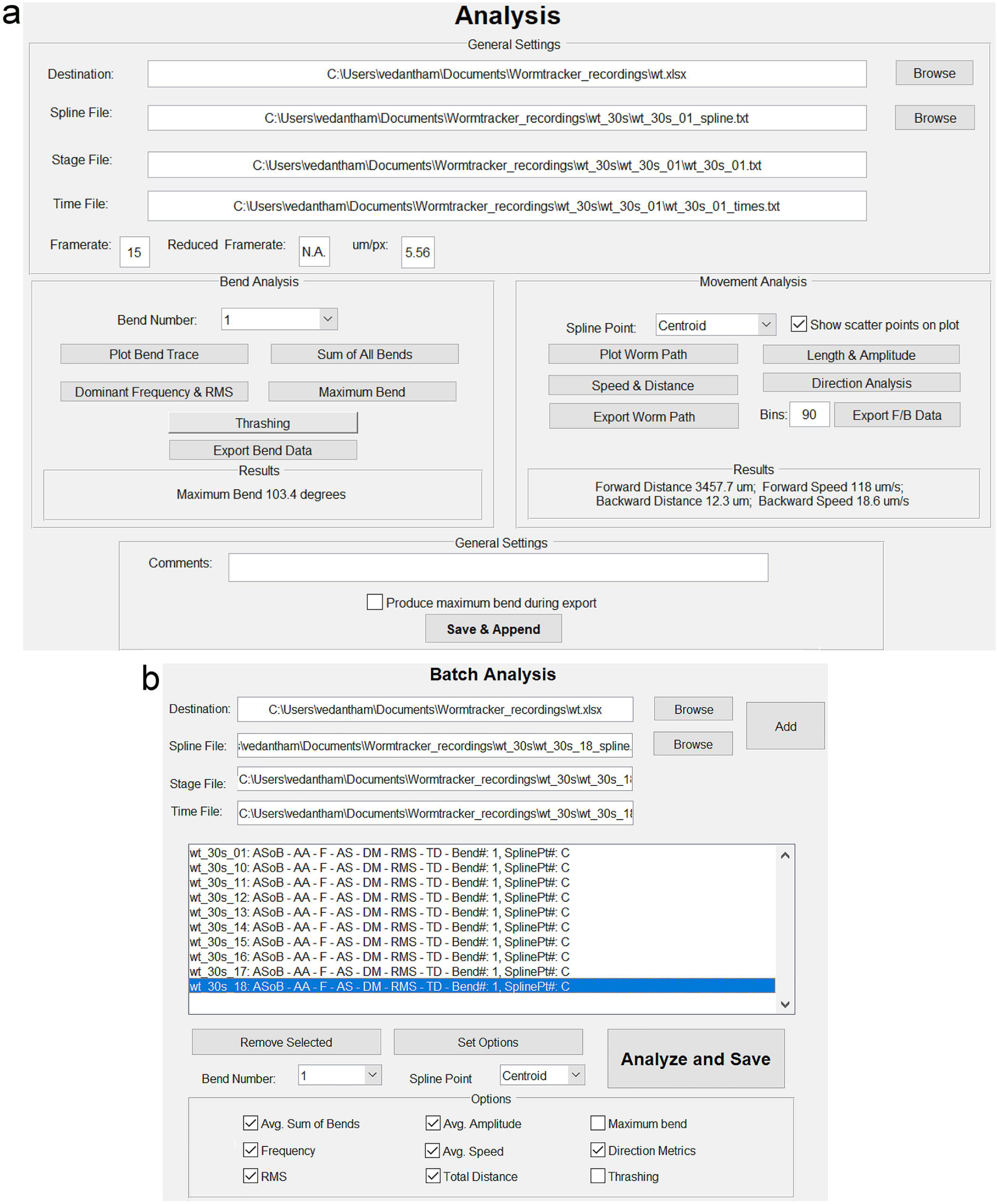
User interfaces of the Analysis and Batch Analysis modules. **a.** User interface of the Analysis module. The module integrates spline fitting results (Spline File), stage movement coordinates (Stage File), frame capturing times (Time File), and the camera calibration factor to perform analysis. It has two sections: Bend Analysis and Movement Analysis. Bend Analysis can plot bend traces, generate bending frequency spectra, display RMS (root mean square) bending angles, and quantify the maximum bending angle for any of 11 selectable bends. It can also display the sum of all RMS bends. Movement Analysis can reconstruct the worm’s travel path, quantify worm length and mean amplitude, calculate average speed and determine both total and net distances traveled based on the centroid or a user-chosen spline point. Additionally, it quantifies distances and average speeds of forward and backward movement and generate a plot of forward/backward speed over time. Users can append results from multiple recordings to the same Excel file (as specified in the Destination field). **b.** User interface of the Batch Analysis module. This module allows users to automatically analyze multiple recordings and save the results in a single Excel file.

Bend Analysis can plot the bend trace and bend frequency spectrum, identify the dominant bend frequency, quantify the RMS and maximum bend amplitudes, and calculate the sum of all RMS bends.

Movement Analysis can reconstruct the worm’s travel path, determine the worm length (L), amplitude (A), and A/L ratio, and quantify the total and net distances travelled, along with the average speed. Additionally, it can quantify forward and backward distances travelled, calculate forward and backward speeds, and generate a graph showing forward and backward speeds over time.

Before using the *Analyze* module, the user is prompted to name an Excel file in a selected folder or subfolder. This ‘master’ Excel file saves all key quantification information, including:

- Analyzed spline file
- Selected spline point
- Selected bend number
- Dominant bend frequency and RMS bend degree of the selected bend
- Sum of all RMS bends
- Worm length (L)
- Worm amplitude (A)
- A/L ratio
- Average speed
- Forward and backward speeds
- Total and net distances travelled
- Forward and backward distances traveled, along with the numbers of frames related to them.

Information from other recordings can be appended to this ‘master’ Excel file, facilitating subsequent statistical analysis.

Additionally, the *Analyze* module can export data for plotting bend traces of all 11 bends, reconstructing the worm’s travel path, and generating a time-series graph of forward and backward speeds as additional Excel files.

#### Batch Analyze

This module (**Figure 10b**) streamlines the analysis of multiple recordings. Users can select quantification parameters for a specific bend number and a designated spline point.

#### Curve Analyzer

This module (**Figure 11**) generates curvature values by fitting the spline with circles, which are displayed as solid (ventral), dashed (dorsal), or dotted (undifferentiated ventral/dorsal) lines. Users can save the curvature values of all curves in an Excel file, which includes the positions (normalized by body length), ventral/dorsal orientation information (if available), and sequential frame numbers.

**Figure 11.**
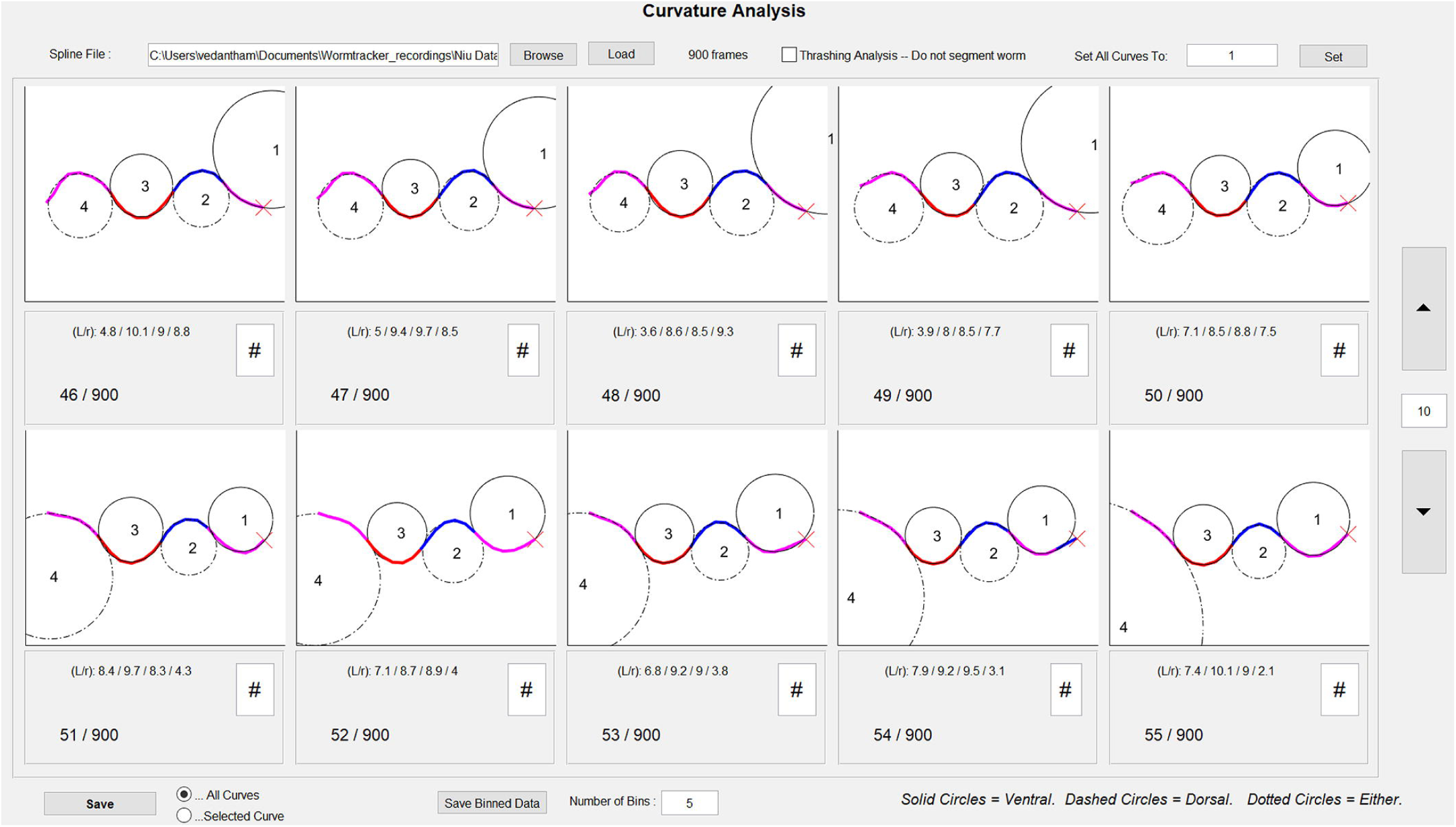
User interface of the Curve Analyzer. The spline of each frame is fitted by circles: solid circles for ventral curvatures, dashed circles for dorsal curvatures, and dotted circles for undifferentiated ventral/dorsal curvatures. The curvature values can be saved either as raw quantified data or as binned data (divided into equal segments of body length) in Excel files.

Since the software can detect even very small curvatures, users may want to exclude these from the analysis. This can be done by filtering out curvature values below a specific threshold using the sorting and editing functions in the Excel file, and then reassigning the remaining curvatures as V1, D1, V2, D2, and so on. Statistical comparisons using the sorted data may make it easier to detect curvature differences between wild-type and mutant animals compared to using the automatically assigned curvature values.

Another option is to save binned curvature data, where the bin size is a fraction of the worm length. The default number of bins is 5. For example, if a mutant differs from wild type primarily in head curvatures, data from Bin 1 (and possibly Bin 2) may be more revealing of the difference.

A third option is to save the curvature values of a specific curve, such as Curve 1 (by selecting “Set All Curves to 1”), to an Excel file. Since the positions and curvature values of the automatically defined Curve 1 can vary greatly from frame to frame, users may choose to use the curvature values of other curves as surrogates for Curve 1 in some frames for the analysis. However, this approach is not recommended for general use because it can be subjective and time-consuming.

#### SleepTracker

SleepTracker offers two user interfaces: Sleep Recorder (**Figure 12a**) and Sleep Analyzer (**Figure 12b**). Sleep Recorder allow users to record the locomotion activity of worms, with customizable exposure time, inter-frame interval, and total recording duration. Up to four worms can be recorded simultaneously using the PDMS membrane with circular openings (300 µm) designed by our team (**Appendix A**). Sleep Analyzer quantifies worm locomotor activity over time, with the exported data used for determining the total sleep duration as well as the number and duration of active events during sleep. Active events during sleep can be detected using the Quick Peaks gadget of the Origin software (OriginLab Corporation, Northampton, MA, USA) based on a threshold of a ≥10 μm difference in centroid location between consecutive images. The duration of motionless sleep can be determined by subtracting the summed duration of active events from the total sleep duration. These methods for quantifying sleep duration and active events during sleep were applied in our study investigating the effects of melatonin and the Slo1 potassium channel on sleep [37].

**Figure 12.**
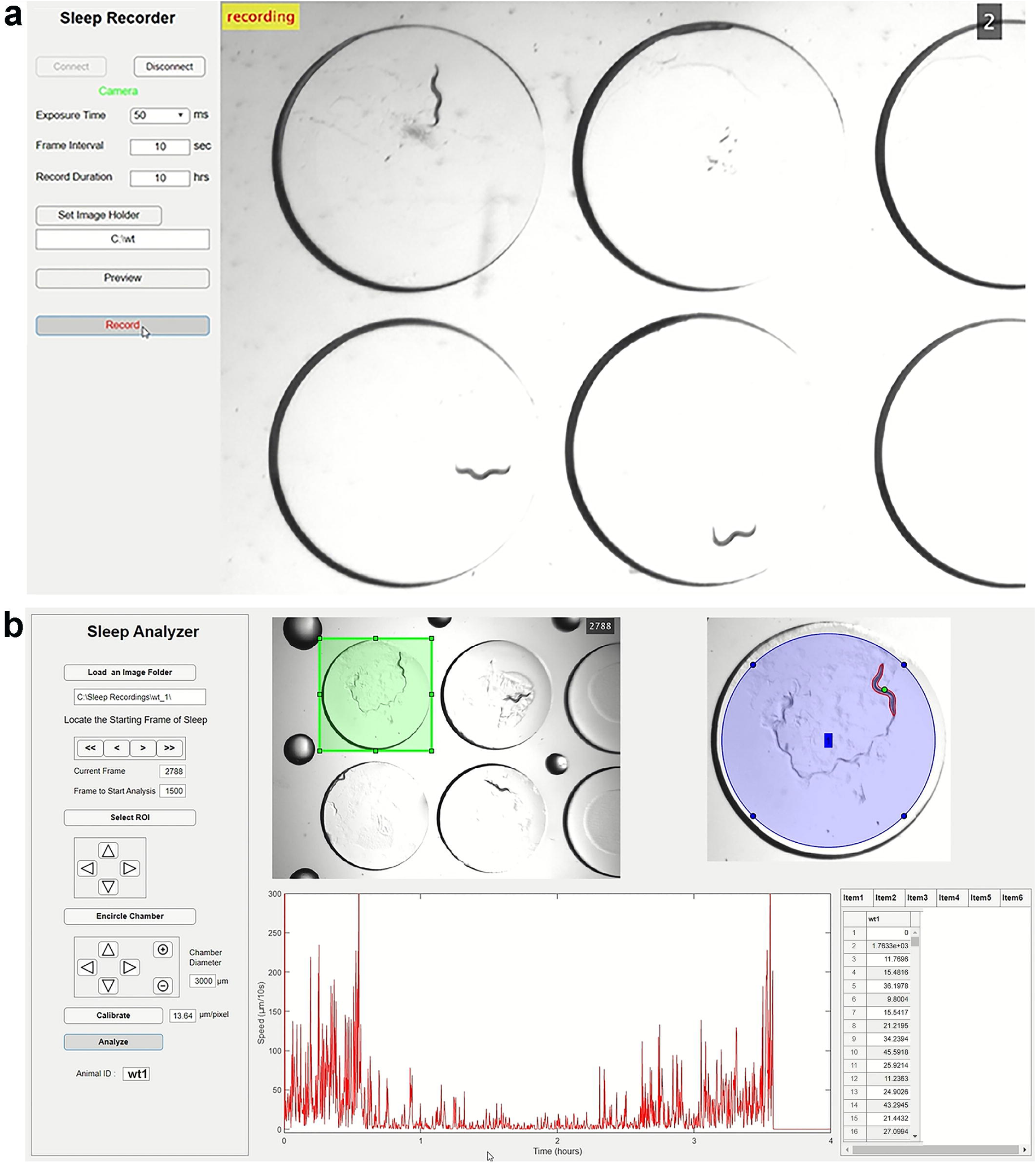
User Interfaces of Sleep Recorder and Sleep Analyzer. **a.** Sleep Recorder allows simultaneous recording of the locomotor activity of four worms using the displayed recording chamber. Users can specify the exposure time, inter-frame interval, and record duration, and save the captured images to a user-specified folder. **b.** Sleep Analyzer quantifies worm locomotor activity over time and displays an actogram of the worm’s movement. The results can be saved as an Excel file for quantifying total sleep duration, as well as the number and duration of active events during sleep.

#### AP Analyzer

AP Analyzer (**Figure 13**) is used to quantify key metrics related to action potentials (APs), including threshold, amplitude, duration at 50% amplitude (APD50), rise and decay times, maximum and minimum slopes, and rise and decay slopes. Additionally, the analyzer measures the afterhyperpolarization (AHP) level and resting membrane potential (RMP), and generates an averaged AP trace along with an AP voltage phase plot.

**Figure 13.**
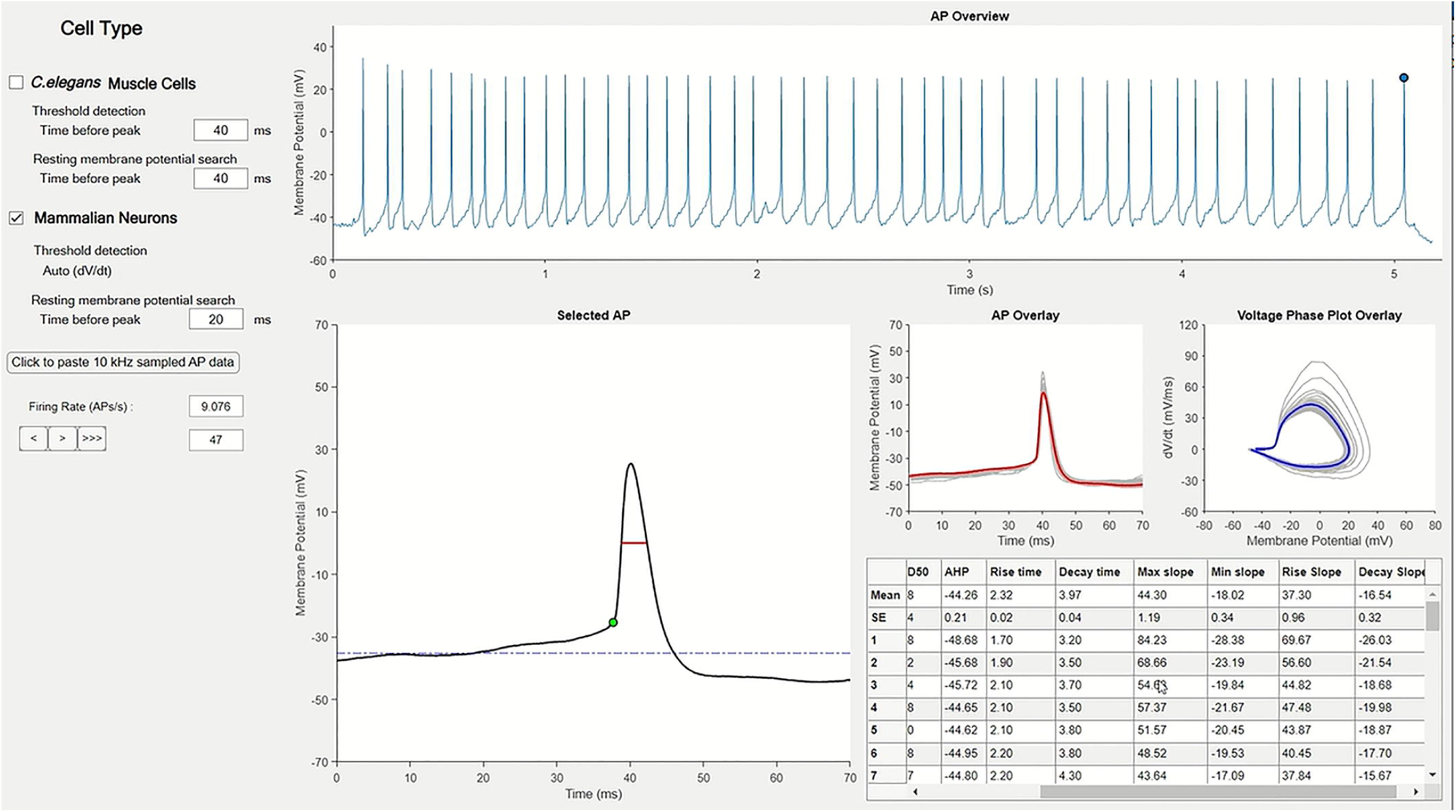
User Interface of AP Analyzer. Users can important current-clamp data acquired at 10 kHz into this module to quantify action potential (AP) metrics and resting membrane potential (RMP). Select “Mammalian Neurons” if the APs have a clear pre-upstroke inflection point, or “*C. elegans* Muscle Cells” if they do not. For APs without a pre-upstroke inflection point, the membrane potential at a user-specified pre-AP peak time is quantified as the threshold. For both mammalian neurons and *C. elegans* muscle cells, RMP is calculated as the average membrane potential over 20 ms before a user-specified pre-AP peak time, which is typically a longer duration than that used for quantifying the AP threshold. The software displays all APs (AP Overview), the AP being analyzed (Selected AP), individual and averaged AP waveforms (AP Overlay), individual and averaged voltage phase plots (Voltage Phase Plot Overlay). It also shows the quantified results, including RMP, AP threshold, AP amplitude, APD50 (AP duration at 50% amplitude), afterhyperpolarization (AHP) level, AP rise and decay times, and maximum and minimum AP slopes. An Excel file can be saved with the quantifying results for individual APs and their averages, along with the data used to generate the averaged AP waveform and AP voltage phase plot.

The RMP is determined by quantifying the membrane potential at a user-defined pre-AP peak time. The AP threshold can be determined using two different approaches, depending on the cell type selected. For APs like those in mammalian neurons, the threshold is determined by identifying the inflection point before the AP upstroke. For APs like those in *C. elegans* body-wall muscle cells, the threshold is defined as the membrane voltage at a user-defined pre-AP peak time. The quantification results are displayed in a table and can be exported to an Excel file, which also includes the data used to generate the averaged AP trace and the AP voltage phase plot.

### EXAMPLES of ANALYSIS

We compared between wild type and several mutant strains in locomotor and bending properties to illustrate various functionalities of *WormTracker*. Several *Unc* (*unc*oordinated) mutants, including *unc-7(e5)*, *unc-9(fc16)*, *unc-8(e1069)*, and *unc-58(n495)*, exhibited significant differences in bending angles, dominant bending frequency, forward and backward speeds compared to wild type (**Figure 14a**). One mutant of a calcium-binding protein, *zw47*, isolated in our genetic screen displayed a significant increase in the body curvature (**Figure 14b**), and a of *lgc-46(ok2949)* spent more time on forward locomotion but less time on backward locomotion, and moved faster in the forward direction but slower in the backward direction compared to wild type (**Figure 14c**).

**Figure 14.**
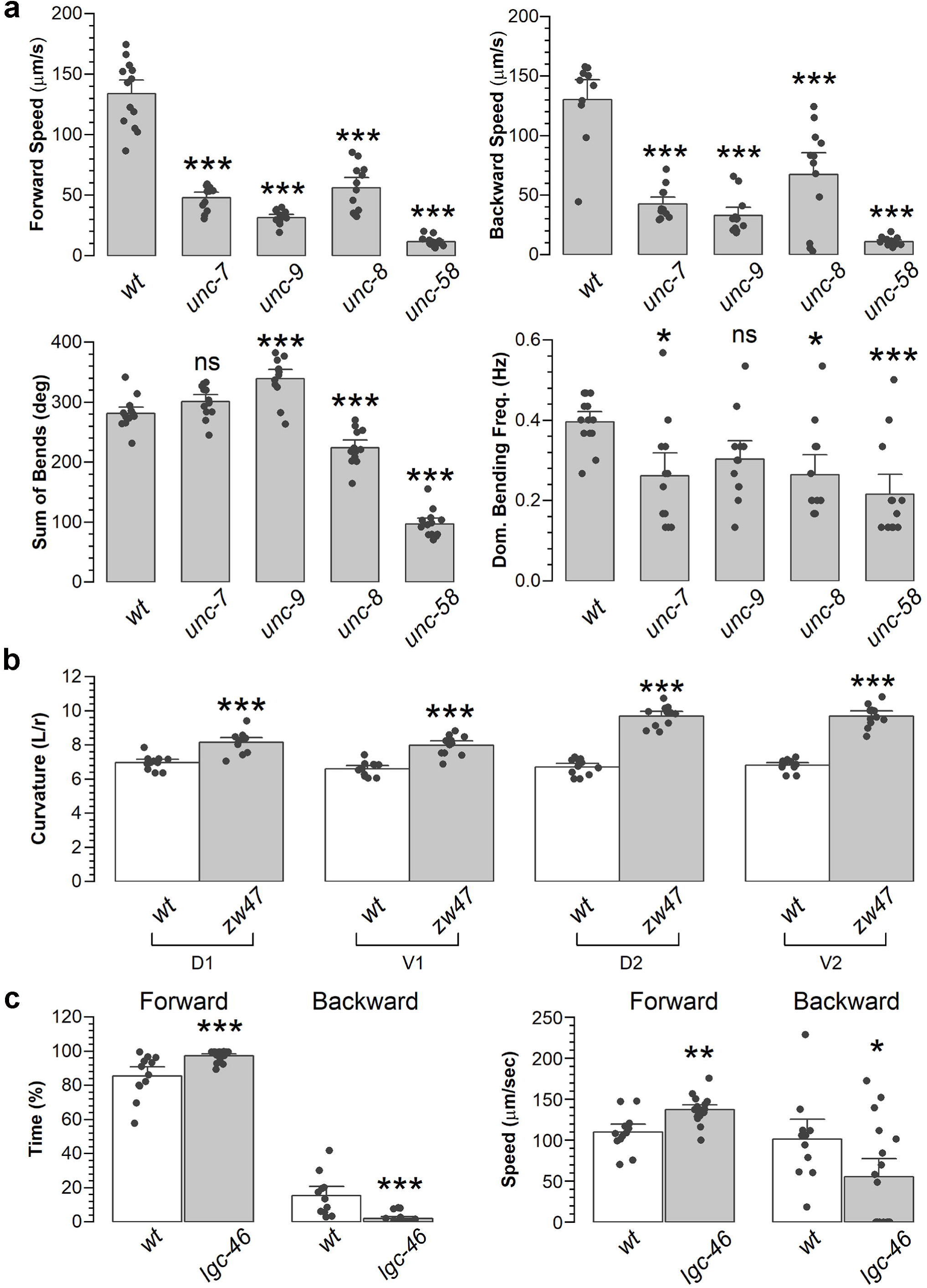
Results of sample worm strains demonstrating functionality of the WormTracker. **a.** Loss-of-function mutations of *unc-7* (innexin), *unc-9* (innexin), *unc-8* (degenerin/epithelial sodium channel), and *unc-58* (potassium channel) result in reduced locomotion speed and abnormal body bending properties compared to wild type (*wt*). The mutant strains analyzed were *unc-7(e5)*, *unc-9(fc16)*, *unc-8(e1069)*, and *unc-58*(n495). **b.** *zw47*, a mutant of a calcium binding protein, displays increased body curvatures compared to *wt*. *Left:* Diagram illustrating the V1, V2, D1, and D2 curvatures. *Right:* Statistical comparisons of body curvatures between *wt* and *zw47*. If the curvature value of the original V1 or D1 was ≥ 2.5, it is excluded from the analysis, with the following V2 or D2 being treated as V1 or D1 in the analysis. **c.** Loss-of-function mutations of *lgc-46* (an acetylcholine receptor) [38, 39], inhibits backward locomotion but enhances forward locomotion compared to *wt*. The mutant analyzed was *lgc-46(ok2949)*. The single, double, and triple asterisks indicate statistically significant differences compared to *wt* at “*p* < 0.05”, “*p* < 0.01”, and “*p* < 0.001”, respectively, while “ns” indicates no significant difference compared to *wt* based on one-way *ANOVA* with Tukey’s post hoc test (**a**) or un-paired *t*-test (**b**, **c**).

## DISCUSSION

We have developed a comprehensive software and hardware system designed to quantify worm locomotor behavior and bending properties, identify sleep episodes and quantify their durations in freely moving worms, and determine AP properties in *C. elegans*. To our knowledge, this is the only open-source system that combines these functionalities.

*WormTracker* is both powerful and user friendly, capable of quantifying all locomotor and bending metrics typically needed by *C. elegans* researchers. The system can distinguish between the worm’s ventral and dorsal sides reliably, track a single worm over extended periods by adjusting stage positions between frames, and integrate external devices via TTL signals.

One limitation of *WormTracker* is that images of coiled worms sometimes cannot be automatically fitted to the spline, requiring user-assisted semi-automatic methods. However, this generally does not present a significant problem, as typical recordings for quantifying locomotion and body bending metrics last only 30 seconds to a few minutes.

*SleepTracker* offers an advantage over methods that confine worms within microfluidic devices by quantifying sleep behavior quickly and accurately in freely moving worms. Recordings must be discarded if the worm contacted the edge of the recording chamber, as this complicates automatic binary image generation. However, this is generally not a major issue, as most animals (70-80%) remain near the center of the chamber due to the restriction of food source (OP50 bacteria) to the center.

*AP Analyzer* is designed to work with current-clamp data acquired at a 10 kHz sampling rate, which is sufficient for accurately capturing AP waveforms. For APs with a clear pre-upstroke inflection point, the software automatically detects the AP threshold, enabling accurate quantification of AP metrics. This method is more efficient and objective than the user-assisted manual threshold selection method used in *ClampFit*, which does not account for the effects of baseline fluctuations on measured AP thresholds. *ClampFit* is a module of *pClamp*, the software widely used by electrophysiological labs for data acquisition and analysis. While *AP Analyzer* offers an advantage over *ClampFit* in AP threshold detection, its automatic detection method is not applicable to typical APs in *C. elegans* body-wall muscle cells due to the absence of a pre-upstroke inflection point. In these cases, our alternative approach of defining the threshold from a specific time before the AP peak ensures consistency across APs, though it remains subjective.

*AP Analyzer* also measures the RMP by calculating the average membrane potential over 20 ms before a specific pre-AP peak time. However, the measured RMP can differ significantly from the true RMP when inter-spike intervals are short. We recommend that users, whenever possible, measure the RMP from a segment of the recording without APs to obtain a more accurate measurement.

We hope that many laboratories will find *Track-A-Worm 2.0* to be a valuable tool for their research. As an open-source system, users can contribute to improving its functionality by providing feedback and refining software codes over time.

## Acknowledgements

This work was supported by NIH R01NS109388 (Z. W.) and R01MH085927 (Z.W.). We thank the Caenorhabditis Genetic Center for mutant strains.

## REFERENCES

1. Mori I. Genetics of chemotaxis and thermotaxis in the nematode Caenorhabditis elegans. Annu Rev Genet. 1999;33:399–422. 10.1146/annurev.genet.33.1.399

2. Goodman MB. Mechanosensation. WormBook. 2006;10.1895/wormbook.1.62.11-14. 10.1895/wormbook.1.62.1

3. Bargmann CI. Chemosensation in *C. elegans*. WormBook. 2006;10.1895/wormbook.1.123.11-29. 10.1895/wormbook.1.123.1

4. Ardiel EL, Rankin CH. C. elegans: social interactions in a “nonsocial” animal. Adv Genet. 2009;68:1–22. 10.1016/S0065-2660(09)68001-9

5. Giles AC, Rose JK, Rankin CH. Investigations of learning and memory in Caenorhabditis elegans. Int Rev Neurobiol. 2006;69:37–71. 10.1016/S0074-7742(05)69002-2

6. Ardiel EL, Rankin CH. An elegant mind: learning and memory in Caenorhabditis elegans. Learn Mem. 2010;17:191–201. 10.1101/lm.960510

7. Nichols ALA, Eichler T, Latham R, Zimmer M. A global brain state underlies C. elegans sleep behavior. Science. 2017;356:10.1126/science.aam6851

8. Barr MM, Garcia LR. Male mating behavior. WormBook. 2006;10.1895/wormbook.1.78.11-11. 10.1895/wormbook.1.78.1

9. Liu P, Ge Q, Chen B, Salkoff L, Kotlikoff MI, Wang ZW. Genetic dissection of ion currents underlying all-or-none action potentials in *C. elegans* body-wall muscle cells. J Physiol. 2011;589:101–17. 10.1113/jphysiol.2010.200683

10. Gao S, Zhen M. Action potentials drive body wall muscle contractions in *Caenorhabditis elegans*. Proc Natl Acad Sci U S A. 2011;108:2557–62. 10.1073/pnas.1012346108

11. Liu Q, Kidd PB, Dobosiewicz M, Bargmann CI. C. elegans AWA Olfactory Neurons Fire Calcium-Mediated All-or-None Action Potentials. Cell. 2018;175:57–70 e17. 10.1016/j.cell.2018.08.018

12. Jiang J, Su Y, Zhang R, Li H, Tao L, Liu Q. C. elegans enteric motor neurons fire synchronized action potentials underlying the defecation motor program. Nat Commun. 2022;13:2783. 10.1038/s41467-022-30452-y

13. Consortium CeS. Genome sequence of the nematode C. elegans: a platform for investigating biology. Science. 1998;282:2012–8. 10.1126/science.282.5396.2012

14. Nurk S, Koren S, Rhie A, Rautiainen M, Bzikadze AV, Mikheenko A et al. The complete sequence of a human genome. Science. 2022;376:44–53. 10.1126/science.abj6987

15. Lander ES, Linton LM, Birren B, Nusbaum C, Zody MC, Baldwin J et al. Initial sequencing and analysis of the human genome. Nature. 2001;409:860–921. 10.1038/35057062

16. Lai CH, Chou CY, Ch’ang LY, Liu CS, Lin W. Identification of novel human genes evolutionarily conserved in Caenorhabditis elegans by comparative proteomics. Genome Res. 2000;10:703–13. 10.1101/gr.10.5.703

17. Javer A, Ripoll-Sanchez L, Brown AEX. Powerful and interpretable behavioural features for quantitative phenotyping of Caenorhabditis elegans. Philos Trans R Soc Lond B Biol Sci. 2018;373:10.1098/rstb.2017.0375

18. Javer A, Currie M, Lee CW, Hokanson J, Li K, Martineau CN et al. An open-source platform for analyzing and sharing worm-behavior data. Nat Methods. 2018;15:645–6. 10.1038/s41592-018-0112-1

19. Puchalt JC, Gonzalez-Rojo JF, Gomez-Escribano AP, Vazquez-Manrique RP, Sanchez-Salmeron AJ. Multiview motion tracking based on a cartesian robot to monitor Caenorhabditis elegans in standard Petri dishes. Sci Rep. 2022;12:1767. 10.1038/s41598-022-05823-6

20. Barlow IL, Feriani L, Minga E, McDermott-Rouse A, O’Brien TJ, Liu Z et al. Megapixel camera arrays enable high-resolution animal tracking in multiwell plates. Commun Biol. 2022;5:253. 10.1038/s42003-022-03206-1

21. Bonnard E, Liu J, Zjacic N, Alvarez L, Scholz M. Automatically tracking feeding behavior in populations of foraging C. elegans. Elife. 2022;11:10.7554/eLife.77252

22. Machino K, Link CD, Wang S, Murakami H, Murakami S. A semi-automated motion-tracking analysis of locomotion speed in the C. elegans transgenics overexpressing beta-amyloid in neurons. Front Genet. 2014;5:202. 10.3389/fgene.2014.00202

23. Feng Z, Cronin CJ, Wittig JH, Jr., Sternberg PW, Schafer WR. An imaging system for standardized quantitative analysis of C. elegans behavior. BMC Bioinformatics. 2004;5:115. 10.1186/1471-2105-5-115

24. Tsibidis GD, Tavernarakis N. Nemo: a computational tool for analyzing nematode locomotion. BMC Neurosci. 2007;8:86. 10.1186/1471-2202-8-86

25. Swierczek NA, Giles AC, Rankin CH, Kerr RA. High-throughput behavioral analysis in C. elegans. Nat Methods. 2011;8:592–8. 10.1038/nmeth.1625

26. Ramot D, Johnson BE, Berry TL, Jr., Carnell L, Goodman MB. The Parallel Worm Tracker: a platform for measuring average speed and drug-induced paralysis in nematodes. PLoS One. 2008;3:e2208. 10.1371/journal.pone.0002208

27. Wen Q, Po MD, Hulme E, Chen S, Liu X, Kwok SW et al. Proprioceptive coupling within motor neurons drives *C. elegans* forward locomotion. Neuron. 2012;76:750–61. 10.1016/j.neuron.2012.08.039

28. Yemini E, Jucikas T, Grundy LJ, Brown AE, Schafer WR. A database of Caenorhabditis elegans behavioral phenotypes. Nat Methods. 2013;10:877–9. 10.1038/nmeth.2560

29. Kwon N, Pyo J, Lee SJ, Je JH. 3-D Worm Tracker for Freely Moving C. elegans. PLoS One. 2013;8:e57484. 10.1371/journal.pone.0057484

30. Wang SJ, Wang ZW. Track-a-worm, an open-source system for quantitative assessment of C. elegans locomotory and bending behavior. PLoS One. 2013;8:e69653. 10.1371/journal.pone.0069653

31. Kwon N, Hwang AB, You YJ, SJ VL, Je JH. Dissection of C. elegans behavioral genetics in 3-D environments. Sci Rep. 2015;5:9564. 10.1038/srep09564

32. Perni M, Challa PK, Kirkegaard JB, Limbocker R, Koopman M, Hardenberg MC et al. Massively parallel C. elegans tracking provides multi-dimensional fingerprints for phenotypic discovery. J Neurosci Methods. 2018;306:57–67. 10.1016/j.jneumeth.2018.02.005

33. Deserno M, Bozek K. WormSwin: Instance segmentation of C. elegans using vision transformer. Sci Rep. 2023;13:11021. 10.1038/s41598-023-38213-7

34. Roussel N, Sprenger J, Tappan SJ, Glaser JR. Robust tracking and quantification of C. elegans body shape and locomotion through coiling, entanglement, and omega bends. Worm. 2014;3:e982437. 10.4161/21624054.2014.982437

35. Leifer AM, Fang-Yen C, Gershow M, Alkema MJ, Samuel AD. Optogenetic manipulation of neural activity in freely moving Caenorhabditis elegans. Nat Methods. 2011;8:147–52. 10.1038/nmeth.1554

36. Chen B, Ge Q, Xia XM, Liu P, Wang SJ, Zhan H et al. A novel auxiliary subunit critical to BK channel function in *Caenorhabditis elegans*. J Neurosci. 2010;30:16651–61.

37. Niu L, Li Y, Zong P, Liu P, Shui Y, Chen B et al. Melatonin promotes sleep by activating the BK channel in *C. elegans*. Proc Natl Acad Sci U S A. 2020;117:25128–37. 10.1073/pnas.2010928117

38. Liu P, Chen B, Mailler R, Wang ZW. Antidromic-rectifying gap junctions amplify chemical transmission at functionally mixed electrical-chemical synapses. Nat Commun. 2017;8:14818. 10.1038/ncomms14818

39. Takayanagi-Kiya S, Zhou K, Jin Y. Release-dependent feedback inhibition by a presynaptically localized ligand-gated anion channel. Elife. 2016;5:10.7554/eLife.21734

