## Supplementary material for "*Track-A-Worm 2.0*: A Software Suite for Quantifying Properties of *C. elegans* Locomotion, Bending, Sleep, and Action Potentials": Table 1

**Table 1: *Track-A-Worm 2.0* Hardware Components and Specifications**

| **Component** | **Model or Product #** | **Supplier** | **Product URL** |
|---|---|---|---|
| Motorized Stage | Optiscan™ ES111 | Prior Scientific | https://www.prior.com/product/optiscan-es111 |
| Stage Controller | OptiScan ES11 | Prior Scientific | https://www.prior.com/product/optiscan-es11-controller |
| Universal Specimen Holder | H473 | Prior Scientific | https://www.prior.com/product/h473 |
| Stage Mounting Bracket | H413 | Prior Scientific | N/A |
| Joystick Control Unit | CS1521DP | Prior Scientific | https://www.prior.com/product/cs152dp |
| Monochrome CMOS Camera | Mako G-040B | Allied Vision | https://www.alliedvision.com/en/camera-selector/detail/mako/g-040/ |
| Power Supply for CMOS Camera | 12V, 2A, 8-pin | Allied Vision | https://www.edmundoptics.com/p/allied-vision-12v-2a-8-pin-hirose-power-supply/43032/# |
| myDAQ University Kit | 781326-01 | National Instruments | https://www.ni.com/en-us/shop/model/mydaq-university-kit.html |
| Fluorescence Stereomicroscope | M165 FC | Leica | https://www.leica-microsystems.com/products/light-microscopes/stereo-microscopes/p/leica-m165-fc/ |
| Hoya Colored Glass Longpass Filters | Hoya Y52 (520nm), 12.5mm Dia., 1mm Thick | Hoya Corporation | https://www.edmundoptics.com/p/hoya-y52-520nm-125mm-dia-1mm-thick-colored-glass-longpass-filter/46497/ |
| Monochrome CMOS Camera | DMK 37BUX273 | The Imaging Source | https://www.theimagingsource.com/en-us/product/industrial/37u/dmk37bux273/ |
