## Appendix A for "*Track-A-Worm 2.0*: A Software Suite for Quantifying Properties of *C. elegans* Locomotion, Bending, Sleep, and Action Potentials"

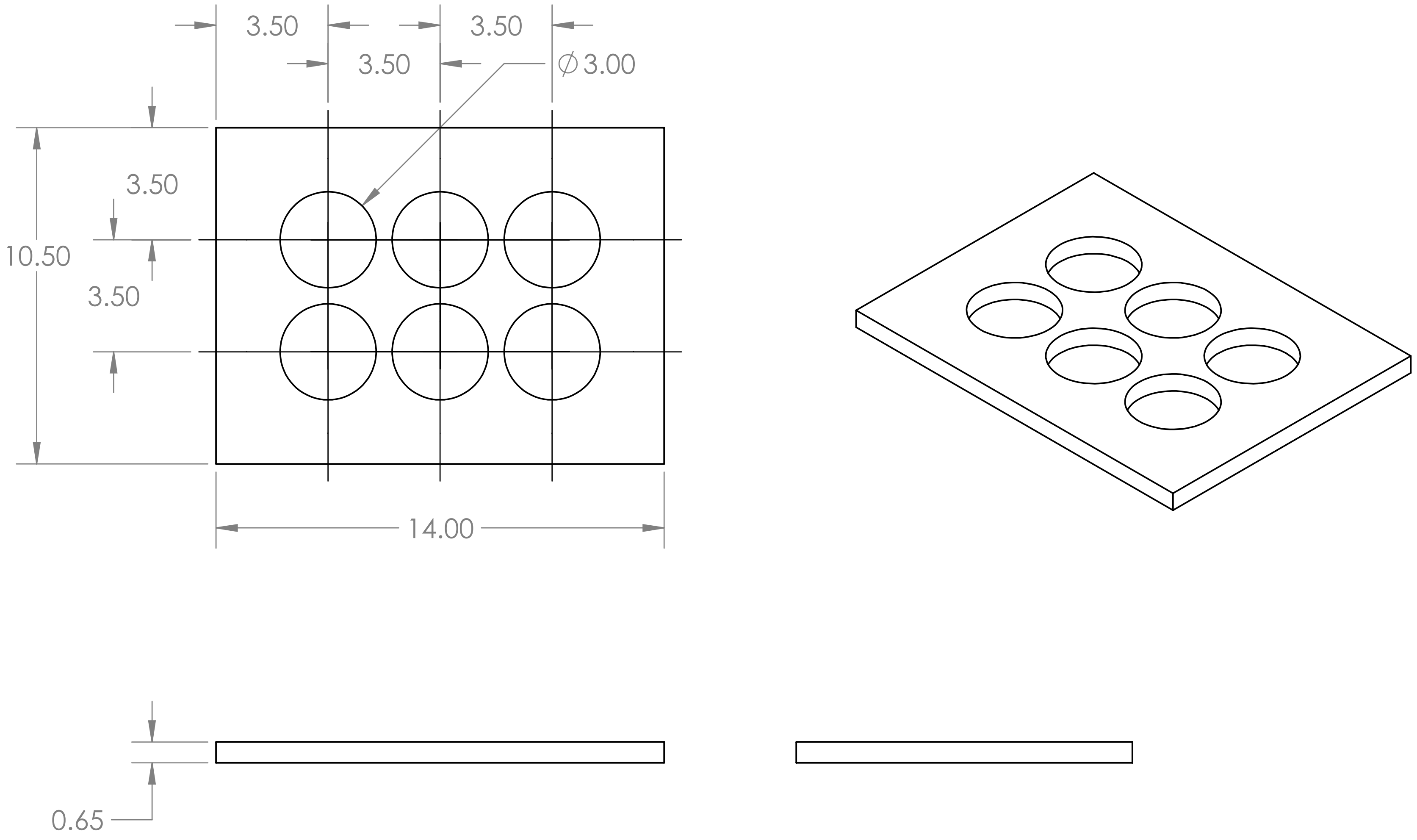

| | |
|---|---|
| NEXT ASSY | USED ON |
| APPLICATION | |

UNLESS OTHERWISE SPECIFIED:

DIMENSIONS ARE IN INCHES
TOLERANCES:
FRACTIONAL ±
ANGULAR: MACH ± BEND ±
TWO PLACE DECIMAL ±
THREE PLACE DECIMAL ±

INTERPRET GEOMETRIC TOLERANCING PER:

MATERIAL

FINISH

DO NOT SCALE DRAWING

| | NAME | DATE |
|---|---|---|
| DRAWN | | |
| CHECKED | | |
| ENG APPR. | | |
| MFG APPR. | | |
| Q.A. | | |

COMMENTS:

TITLE:

| SIZE | DWG. NO. | REV |
|---|---|---|
| A | SixHoleChamber | |

SCALE: 5:1 WEIGHT: SHEET 1 OF 1

**PROPRIETARY AND CONFIDENTIAL**

THE INFORMATION CONTAINED IN THIS DRAWING IS THE SOLE PROPERTY OF <INSERT COMPANY NAME HERE>. ANY REPRODUCTION IN PART OR AS A WHOLE WITHOUT THE WRITTEN PERMISSION OF <INSERT COMPANY NAME HERE> IS PROHIBITED.
